# TiTUS: Sampling and Summarizing Transmission Trees with Multi-strain Infections

**DOI:** 10.1101/2020.03.17.996041

**Authors:** Palash Sashittal, Mohammed El-Kebir

**Affiliations:** Department of Aerospace Engineering, University of Illinois at Urbana-Champaign, Urbama, IL 61801, USA; Department of Computer Science, University of Illinois at Urbana-Champaign, Urbama, IL 61801, USA

## Abstract

**Motivation:** The combination of genomic and epidemiological data hold the potential to enable accurate pathogen transmission history inference. However, the inference of outbreak transmission histories remains challenging due to various factors such as within-host pathogen diversity and multi-strain infections. Current computational methods ignore within-host diversity and/or multi-strain infections, often failing to accurately infer the transmission history. Thus, there is a need for efficient computational methods for transmission tree inference that accommodate the complexities of real data.

**Results:** We formulate the Direct Transmission Inference (DTI) problem for inferring transmission trees that support multi-strain infections given a timed phylogeny and additional epidemiological data. We establish hardness for the decision and counting version of the DTI problem. We introduce TiTUS, a method that uses SATISFIABILITY to almost uniformly sample from the space of transmission trees. We introduce criteria that prioritizes parsimonious transmission trees that we subsequently summarize using a novel consensus tree approach. We demonstrate TiTUS’s ability to accurately reconstruct transmission trees on simulated data as well as a documented HIV transmission chain.

**Availability:** https://github.com/elkebir-group/TiTUS

**Supplementary information:** Supplementary data are available at *Bioinformatics* online.

## 1 Introduction

With the advent of cheaper and more powerful sequencing methods, molecular epidemiology has become an indispensable tool for inference of transmission histories of infectious disease outbreaks. Genomic data of pathogen isolates collected from infected hosts is used to assist with the identification of unknown infection sources and transmission chains. Intensive field work generates crucial epidemiological data that provides addition information such as contact history between patients and exposure times of the patients to sources of infection. Methods that can efficiently use genomic and epidemiological data together for accurate inference of transmission history of outbreaks are the key to real-time outbreak management and devising public health policies and disease control strategies for future outbreaks (Dellicour *et al*., 2018).

There are several challenges that hinder the accurate inference of the transmission history of an outbreak. Phylogeny estimation of the pathogen isolates reveals the evolutionary history of the pathogen during the outbreak. However, due to within-host diversity of the pathogen, branching events in the phylogeny do not correspond to the transmission events during the outbreak (Romero-Severson *et al*., 2014). Phylogeny-based methods that assume that the transmission events coincide with the branching events in the phylogeny are therefore not applicable in the context of pathogens with low mutation rates, short incubation times and acute infections (Ypma *et al*., 2011; Harris *et al*., 2010; Leitner *et al*., 1996; Cottam *et al*., 2008).

Another factor that makes outbreak transmission history inference challenging is a *weak transmission bottleneck*, where multiple strains of the pathogen are transmitted from a donor to a recipient through a non-negligibly small inoculum. Due to this, the most recent common ancestor of lineages from the same host need not have arisen in that host. Although large inocula have been observed in a number of diseases (Leonard *et al*., 2017), most of the existing methods for transmission tree inference that account for the within-host diversity do not account for the co-transmission of pathogen strains (Ypma *et al*., 2013; Didelot *et al*., 2014; Hall *et al*., 2015; Didelot *et al*., 2017). That is, these methods assume a *strong transmission bottleneck* where a single strain of the pathogen is transmitted in an infection. A weak transmission bottleneck is considered in SCOTTI (De Maio *et al*., 2016) and BadTrIP (De Maio *et al*., 2018), however they make the simplifying assumption that all the transmissions are independent of each other. SharpTNI (Sashittal and El-Kebir, 2019) considers the weak transmission bottleneck without this assumption under a parsimony based framework for a known phylogeny. However, SharpTNI may yield transmission histories that cannot be represented by a tree due to multiple infections of a single host from distinct donors. Such super-infection are unlikely for pathogens where infected hosts acquire immunity towards further infections of the pathogen (Whittle *et al*., 1999; Wearing and Rohani, 2009), thus restricting the transmissions history to a tree.

The contributions of this paper are three-fold. First, we consider the problem of counting and sampling uniformly from the set of possible transmission trees for a known phylogeny and epidemiological data. In previous works, this problem is considered by Kenah *et al*. (2016) when the order of infections during the outbreak is completely known and by Hall and Colijn (2019) under the strong transmission bottleneck constraint. In this work, we relax both these constraints and propose a method TiTUS that approximately counts and almost uniformly samples the transmission trees under a weak transmission bottleneck for a given timed phylogeny (Fig. 1). We prove the hardness of the decision and counting versions of this problem and demonstrate the efficiency and accuracy of TiTUS on simulated data. Second, we present a robust criteria for ranking or prioritizing the uniformly sampled candidate transmission trees. In addition to the simulated data, we demonstrate the performance of the selection criteria on an HIV outbreak with a known transmission chain (Vrancken *et al*., 2014). Third, in practice, the underlying phylogeny has some uncertainty and there can be multiple candidates for the transmission tree for a given phylogeny. It is therefore desirable to have an efficient method to summarize the solution space of transmission trees that are consistent with the genetic and epidemiological data. To this end, we propose a consensus-based method that provides the mean transmission tree for a set of candidate solutions while accounting for the number of distinct strains transmitted in each infection event.

**Fig. 1:**
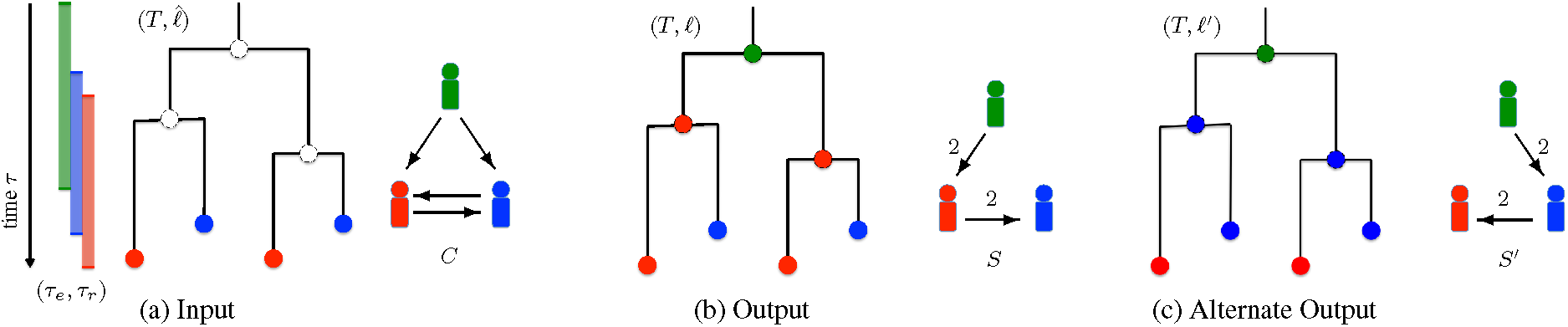
Overview of the Direct Transmission Inference (DTI) problem. (a) The input of the problem consists of a timed phylogeny *T* that captures the evolutionary history of the pathogen during the course of the outbreak. Each leaf of *T* corresponds to a sample collected for an individual host and is thus labeled using 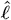 (indicated by colors). The entry and removal times [*τ*_*e*_(*s*), *τ*_*r*_ (*s*)] for each host *s* is also included in the input. (b) Our aim is to label the internal vertices of *T* with *ℓ* such that the resulting transmission edges form a transmission tree *S* (as shown in Fig. 1b). Each edge (*s, t*) of *S* is weighted by the number of transmission edges from host *s* to host *t* given by the vertex labeling *ℓ*. (c) An alternative solution to the given DTI instance. It is easy to see that no solution exists under the strong bottleneck constraint whereas under the weak transmission bottleneck there are multiple solutions. All the feasible vertex labelings are shown in Fig. S7.

## 2 Preliminaries

To state the problems we consider in this manuscript, we start by introducing the required concepts and notation. Let *T* be a rooted tree with vertex set *V*(*T*) and edge set *E*(*T*). The set of leaves of the tree is given by *L*(*T*). The root of the tree is denoted by *r*(*T*). We denote the children of a vertex *u* by *δ*_*T*_ (*u*). We write *u* ⪯_*T*_ *v* if vertex *u* is ancestral to vertex *v, i*.*e*. vertex *u* is present on the unique path from *r*(*T*) to vertex *v*. Note that ⪯_*T*_ is reflexive, *i*.*e*. it holds that *u* ⪯_*T*_ *u* for all vertices *u*. We denote the set of *m* distinct hosts in the outbreak by Σ. In a phylogeographical setting, the set Σ corresponds to *m* distinct geographical locations.

The evolutionary of all strains of a pathogen in an outbreak is modeled by a timed phylogeny, which we define as follows.

### Definition 1.

A *timed phylogeny T* is a rooted tree whose vertices are labeled by time-stamps *τ* : *V* (*T*) → ℝ^≥0^ such that *τ* (*u*) ≤ *τ* (*v*) for all pairs *u, v* of vertices where *u* ⪯_*T*_ *v*.

Thus, as we can see in the above definition, time moves forward when traversing down a timed phylogeny *T* starting from the root *r*(*T*). It is important to note that the leaves of a timed phylogeny *T* may occur at distinct time-stamps, *i*.*e. T* is not necessarily ultrametric.

Each leaf of a timed phylogeny *T* corresponds to a strain of pathogen that was collected during the outbreak. As such, we know the host from which each individual strain was isolated. This is captured by a leaf labeling, *i*.*e*. a labeling of the leaves of *T* by hosts Σ.

### Definition 2.

A *leaf labeling* of a timed phylogeny *T* is a function 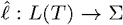, assigning a host 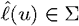 to each leaf vertex *u* ∈ *L*(*T*).

While we know the host 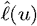 from which each individual leaf *u* of *T* was sampled, we do not know the hosts of the internal vertices, which correspond to unsampled, ancestral strains. Here, our goal is to determine the hosts in which these ancestral strains reside.Mathematically, we wish to construct a *vertex labeling ℓ* : *V* (*T*) → Σ, such 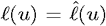 for all leaves *u* ∈ *L*(*T*). Given a vertex labeling *ℓ*, each internal vertex *u* of *T* thus corresponds to a strain residing within host *ℓ*(*u*) at time *τ* (*u*).

In addition to the evolutionary history of all strains in the outbreak, a timed phylogeny *T* combined with a vertex labeling *ℓ* gives us information about the transmission history of the outbreak. Transmissions of strains from one host to another correspond to edges (*u, v*) of *T* labeled by distinct hosts *ℓ*(*u*) ≠ *ℓ*(*v*). Formally, we define a *transmission edge* as follows.

### Definition 3.

Given a timed phylogeny *T* and vertex labeling *ℓ*, an edge (*u, v*) of *T* is a *transmission edge* if *ℓ*(*u*) ≠ *ℓ*(*v*).

The vertex labeling that we construct for a given timed phylogeny *T* and leaf labeling 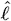, must follow certain constraints for a realistic reconstruction of the transmission history of the pathogen. We will now define these epidemiological constraints.

The first constraint that we introduce is called the *direct transmission* constraint, which imposes the following two restrictions. First, the outbreak begins with a single infected host. We call this initial host the *root host* and it labels the root node *r*(*T*) of the timed phylogeny. The *root host* is not infected by any other host and therefore if *s* is the root host, there cannot exist a transmission edge (*u, v*) such that *ℓ*(*u*) ≠ *s* and *ℓ*(*v*) = *s*.Second, the remaining hosts have a unique infector and are thus infected only once in the course of the outbreak. A possible explanation for this phenomenon is diseases where infected hosts acquire immunity towards further infections of the pathogen (Whittle *et al*., 1999; Wearing and Rohani, 2009). Consequently, there cannot exist two distinct transmission edges (*u, v*) and (*u*′, *v*′) such that *ℓ*(*v*) = *ℓ*(*v*′) and *ℓ*(*u*) ≠ *ℓ*(*u*′). However, an infection between any two hosts *s, t* ∈ Σ may involve the transmission of multiple strains at the same time. This is known as a *weak transmission bottleneck*. Since the transmission of strains must occur concurrently, the time intervals corresponding to any two transmission edges between the same pair (*s, t*) of hosts must have an non-empty intersection. More formally, we state the *direct transmission* constraint as follows,

### Definition 4.

For a timed phylogeny *T*, a vertex labeling *ℓ* satisfies the *direct transmission constraint* if (i) there does not exist a transmission edge (*u, v*) such that *ℓ*(*v*) = *ℓ*(*r*(*T*)) and (ii) we have [*τ* (*u*), *τ* (*v*)] ∩ [*τ* (*u*′), *τ* (*v*′)] ≠ ø for any two transmission edges (*u, v*) and (*u*′, *v*′) where *ℓ*(*u*) = *ℓ*(*u*′) and *ℓ*(*v*) = *ℓ*(*v*′).

Under the *direct transmission* constraint, the set of transmission edges induced by the vertex labeling *ℓ* uniquely determines the *transmission tree S*. More formally, the vertex set *V* (*S*) of a transmission tree *S* is the host set Σ, and there is a directed edge from *s* ∈ Σ to *t* ∈ Σ if and only if there exists at least one edge (*u, v*) ∈ *E*(*T*) such that (i) *s* ≠ *t*, (ii) *ℓ*(*u*) = *s* and (iii) *ℓ*(*v*) = *t*. Since every host except the *root host* has a unique infector, the directed edges necessarily form a tree. Each directed edge (*s, t*) ∈ *E*(*S*) is given a weight *w* : *E*(*S*) → ℕ such that *w*(*s, t*) equals the number of transmission edges in *T* from host *s* to *t*. If *w*(*s, t*) = 1 for all edges (*s, t*) ∈ *E*(*S*) then each host is infected due to the transmission of a single pathogen strain. This phenomenon is known as a *strong transmission bottleneck*.

Epidemiological data provide two additional types of information. First, for each host *s* we are given an interval [*τ*_*e*_(*s*), *τ*_*r*_ (*s*)] during which the host was present in the outbreak and susceptible for infection. Specifically, *τ*_*e*_(*s*) ∈ ℝ^≥0^ is the entry time at which host *s* became susceptible for infection, whereas *τ*_*r*_ (*s*) ∈ ℝ^≥0^ is the *removal time* at which the host was removed from the susceptible and infected populations and placed in treatment or recovering.

Second, there can also be documented geographical constraints that prevent transmissions between any given pair of hosts. We account for all such constraints using a *contact map*. A *contact map C* is a directed graph whose vertex set equals the set Σ of hosts. A directed edge (*s, t*) represents a possible infection event from host *s* to host *t*. If any two hosts are not connected in *C* then there can be no infection event between that pair of hosts. It can clearly be seen that (i) the contact map *C* is a subgraph of the interval graph induced by the intervals [*τ*_*e*_(*s*), *τ*_*r*_ (*s*)], ∀*s* ∈ Σ and (ii) the transmission tree *S* is a spanning arborescence of the contact map *C*. Thus, even in the absence of documented contacts between hosts, a contact map is induced by the entry and removal times of the hosts.

## 3 Problem Statement

We focus on inferring the transmission history of an outbreak for a known pathogen phylogeny *T*. In addition, we are given epidemiological data, which include the contact map *C*, entry and removal times [*τ*_*e*_(*s*), *τ*_*r*_ (*s*)] for each host *s* ∈ Σ and assume a direct transmission constraint under a weak transmission bottleneck. This leads to the following decision problem.

### Problem 1

(Direct Transmission Inference (DTI)). Given a timed phylogeny *T* with time-stamps *τ* : *V* (*T*) → ℝ^≥0^, a leaf labeling 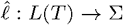, a contact map *C* and entry *τ*_*e*_ : Σ → ℝ^≥0^ and removal times *τ*_*r*_ : Σ → ℝ^≥0^, find a vertex labeling *ℓ* that induces a transmission tree *S* that is a spanning arborescence of *C* and *τ* (*u*) ∈ [*τ*_*e*_(*s*), *τ*_*r*_ (*s*)] for all hosts *s* and vertices *u* where *ℓ*(*u*) = *s*.

An instance of the DTI problem is shown in Fig. 1a shows an instance of the DTI problem along a with a solution vertex labeling *ℓ* and induced transmission tree *S*, where the three hosts are inducated using three colors. Importantly, a DTI problem instance may admit multiple solutions, as shown in Fig. 1b and Fig. 1c. These solutions provide alternative reconstructions of the transmission history, and thus must be taken into consideration in any downstream analysis of the outbreak to devise policy to better manage/prevent future outbreaks. To quantify the number of alternative reconstructions, we formulate the following counting problem.

### Problem 2

(# Direct Transmission Inference (#DTI)). Given a timed phylogeny *T* with time-stamps *τ* : *V* (*T*) → ℝ^≥0^, a leaf labeling 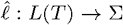, a contact map *C* and entry *τ*_*e*_ : Σ → ℝ^≥0^ and removal times *τ*_*r*_ : Σ → ℝ^≥0^, count the number of vertex labelings *ℓ* that induce a transmission tree *S* that is a spanning arborescence of *C* and *τ* (*u*) ∈ [*τ*_*e*_(*s*), *τ*_*r*_ (*s*)] for all hosts *s* and vertices *u* where *ℓ*(*u*) = *s*.

Let ℒ be the set of all solutions to a given DTI problem instance. Ideally, we would exhaustively enumerate all solutions to the problem instance. However, worst case, the number of solutions scales exponentially with our input. Thus, to obtain a good overview of the solution space ℒ, we need to consider the sampling version of #DTI problem where we wish to uniformly sample the solution space.

In summary, we defined three versions of the DTI problem: a decision, counting and sampling version. In the following, we will consider a previously defined constrained version of the DTI problem as well as a generalization.

### 3.1 Related Transmission Tree Inference Problems

We start by considering a version of the DTI problem with one additional constraint. This additional constraint requires that only one pathogen strain is transmitted to a new host in a transmission event, and is known as a *strong transmission bottleneck*. We refer to this problem as Directed Transmission Inference under Strong Bottleneck (DTI-SB), and denote the space of solutions by ℒ*SB*. This problem was posed by Hall *et al*. (2015).In subsequent work, Hall and Colijn (2019) introduced a polynomial time algorithm to enumerate and uniformly sample from the set ℒ_SB_. Since the DTI-SB only has one additional constraint over the original DTI problem, the solution space of DTI-SB is a proper subset of the solution space of DTI for the same timed phylogeny *T*, leaf labeling 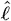 and epidemiological data. More formally, we have ℒ_*SB*_ ⊆ ℒ.

The second problem we consider is a relaxed version of DTI. Specifically, we relax the *direct transmission* constraint for a given instance of DTI. We refer to this problem as rel-DTI and the space of feasible solutions for a given instance by ℒ_REL_. Section 5.2.1 introduces a polynomial time dynamic programming algorithm that enumerates, counts and uniformly samples from the set ℒ_REL_. Since the rel-DTI problem is a relaxation of the DTI problem, we can use the algorithm introduced in Section 5.2.1 to uniformly sample from the solution space of the DTI problem (ℒ). Fig. 2 shows the relation between the solution spaces of the three transmission tree inference problems.

**Fig. 2:**
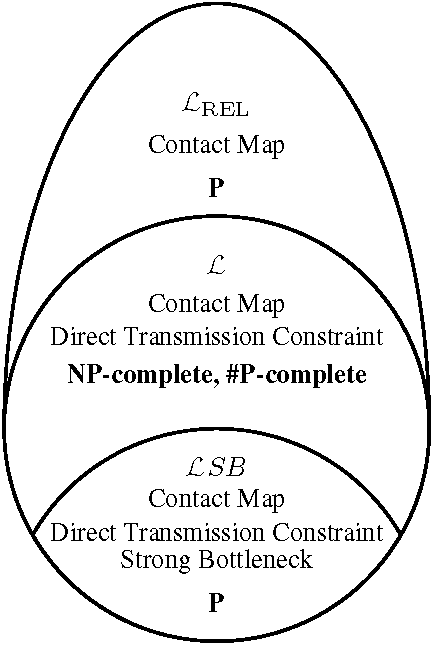
Schematic to compare the solution spaces of transmission trees under different constraints for a known timed phylogeny. We have ℒ_SB_ ⊆ ℒ ⊆ ℒ_REL_. ℒ_SB_ is the solution space of transmission trees with a strong bottleneck that is considered in the work of Hall and Colijn (2019) where they show that counting the solutions and sampling from this solution space can be performed in polynomial time. ℒ is the solution space of DTI which we show to be both NP-complete and #P-complete. Finally, ℒ_REL_ is the relaxed solution space that is used to construct a polynomial time rejection based naive sampling and counting algorithm in Section 5.2.1.

### 3.2 Consensus Tree Problem

For the DTI problem described in the previous section, we start with a given pathogen phylogeny *T*. However, in practice the phylogeny needs to be inferred from genomic sequences of the strains collected from individual hosts Σ. Several methods of phylogeny inference generate either multiple candidates for the phylogeny or a posterior on the solution phylogeny space (Bouckaert *et al*., 2019; Stamatakis, 2014). Moreover, for each given timed phylogeny, we can get multiple solutions to the DTI problem as shown for a representative instance in Fig. 1. Therefore, there is a need for an efficient method to summarize the candidate transmission trees that explain the disease outbreak.

A common method to summarize the solution space of transmission trees is to aggregate the information from the candidate transmission trees to generate a single graph where each edge is weighted by the number of candidate trees that support that edge (De Maio *et al*., 2016; Wymant *et al*., 2017; Didelot *et al*., 2014). This graph rarely represents a single coherent transmission tree among the set of all hosts in the dataset. For this reason, the resulting graph is called a *relationship graph* (Wymant *et al*., 2017) and does not provide crucial information about co-occurrence and mutual exclusivity among edges of the candidate transmission trees.

Another line of method summarizes the set of candidate solutions using one or more consensus trees that best represent the solution space (Jombart *et al*., 2017; Kendall *et al*., 2018). For instance, Jombart *et al*. (2017) apply pairwise distance metrics on the space 𝒮 of transmission trees, not taking into account the number *w*(*s, t*) of transmitted strains between pairs of host (*s, t*). The resulting distance matrix is subsequently embedded into lower dimensional space that the authors then cluster. Finally, each cluster is then assigned a single transmission tree in 𝒮 as its representative (Hall and Colijn, 2019). Kendall *et al*. (2018) follow a similar embedding approach, again not taking the number *w*(*s, t*) of transmission into account. Thus neither method supports a weak transmission bottleneck. To address this limitation, we define the weighted parent-child distance (WPCD) *d*(*S*_1_, *S*_2_) between any two transmission trees *S*_1_ and *S*_2_ as follows.

#### Definition 5.

Let *S*_1_ = (Σ, *E*_1_) with edge labeling *w*_1_ and *S*_2_ = (Σ, *E*_2_) with edge labelings *w*_2_ be two transmission tree on the same vertex set Σ. The *weighted parent-child distance* between the two graphs denoted by *d*(*S*_1_, *S*_2_) is

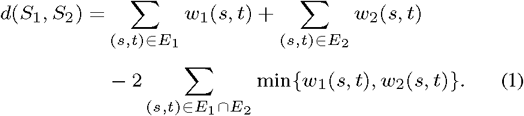

In Appendix A.1.1 we show that this distance function induces a metric in the space 𝒮 of transmission trees. Note that transmission trees *S* and *S*′ that have the same topology but different edge weights *w* and *w*′ will have *d*(*S, S*′) > 0. As a result, WPCD can be used to produce a consensus transmission tree while taking an incomplete transmission bottleneck into account. Under the *strong transmission bottleneck* the *weighted parent-child distance* simplifies to the size of the symmetric difference between the edge sets of the two transmission trees, *i*.*e. d*(*S, S*′) = |*E*′\*E*|+|*E*\*E*′|. This distance is known as the parent-child distance, and has been used to compare tumor phylogenies (Aguse *et al*., 2019; Govek *et al*., 2018). Using WPCD, we define the following consensus tree problem.

#### Problem 3

(Single Consensus Transmission Tree (SCTT)). Given *k* distinct transmission trees 𝒮 = {*S*_1_, …, *S*_*k*_} with edge labelings {*w*_1_, …, *w*_*k*_} find a consensus transmission tree *R* that minimizes 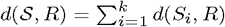.

## 4 Complexity

This section establishes hardness results for the decision and counting versions of the DTI problem.

### Theorem 1.

DTI is NP-complete.

We show the hardness of DTI by reduction from the 1-in-3SAT problem, which is a known NP-complete problem (Karp, 1972). Details are in Appendix A.2.

It is known that the #1-in-3SAT is a #P-complete problem (Creignou and Hermann, 1993). In order to show that the #DTI is also #P-complete, it suffices to show that there exists a polynomial-time reduction from #1-in-3SAT such that the number of solutions is preserved, which we do in Appendix A.2.

### Theorem 2.

#DTI is #P-complete.

Since the decision problem DTI is NP-complete, there does not exist a fully polynomial randomized approximate scheme (FPRAS) for the counting version of DTI unless NP=RP (Jerrum, 2003; Miklós, 2019).

## 5 Methods

This sections describes the methods developed to solve the decision, counting and sampling versions of the DTI problem.

### 5.1 Decision Problem

Since the DTI is NP-complete, we propose to use SATISFIABILITY to solve the decision problem. As such, we construct a Boolean formula *ϕ* for a given DTI instance *T*, 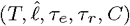, such that there is a bijection between the solutions of the DTI instance and the corresponding SAT instance *ϕ*. Solving the SAT instance will then be equivalent to solving the corresponding DTI problem.

#### Vertex labeling

Decision variables **x** ∈ {0, 1}^*n*×*m*^ encode a vertex labeling, *i*.*e. x*_*i,s*_ = 1 if and only if the node *ℓ*(*v*_*i*_) = *s* and *x*_*i,s*_ = 0 otherwise. We encode uniqueness of the label of each vertex with the following formula.

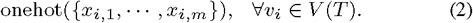

The function onehot(*X*) encodes that exactly one binary variable *x* ∈ *X* is true, which can be accomplished by the following constraint.

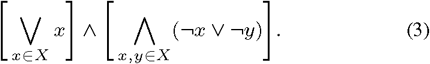

#### Transmission edges

We encode the transmission edges using variables *c*_*s,t*_ with *s, t* ∈ Σ and *s* ≠ *t*. We enforce that *c*_*s,t*_ = 1 if and only if the host *t* is infected by host *s* and *c*_*s,t*_ = 0 otherwise as follows.

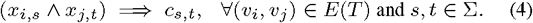

#### Root host

To enforce that the host which labels *r*(*T*) is not infected by any other host, we have

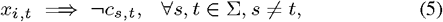

where *v*_*i*_ = *r*(*T*).

#### Direct transmission constraint

We enforce that any host cannot be infected by more than one other host. For each host *s* ∈ Σ we have

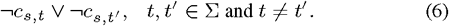

We require that all transmission edges from host *s* to host *t* must have time intervals that overlap. For all edge pair (*v*_*i*_, *v*_*j*_), (*v*_*k*_, *v*_*l*_) that do not have overlapping time intervals, i.e. [*τ* (*v*_*i*_), *τ* (*v*_*j*_)] ∩ [*τ* (*v*_*k*_), *τ* (*v*_*l*_)] = ø, we impose

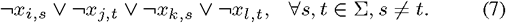

### 5.2 Counting and Sampling Problem

#### 5.2.1 Naive Rejection based Method

For a naive rejection sampling algorithm, we relax the *direct transmission constraint* and uniformly sample vertex labelings for the timed phylogeny *T* such that for all transmission edges (*u, v*) we have (*ℓ*(*u*), *ℓ*(*v*)) ∈ *E*(*C*). As described in Section 3.1, we refer to this as the rel-DTI problem. Let the set of such vertex labelings be ℒ_REL_. Drawing a vertex labeling labeling *ℓ* ∈ ℒ_REL_ uniformly at random from the set ℒ_REL_ can be done in polynomial time, as we describe in Appendix A.3. The sampled vertex labeling labeling *ℓ* is rejected unless it satisfies the *direct transmission constraint*, which can be verified in polynomial time. The probability of success for this rejection based sampling algorithm is 1−(|ℒ|*/*|ℒ_REL_|)^*K*^ after *K* repetitions.

#### 5.2.2 Approximate Counting and Sampling using SAT

Using the SAT formulation shown in Section 5.1, we may use ApproxMC (Chakraborty *et al*., 2013; Soos and Meel, 2019) to approximate |ℒ| and UniGen (Chakraborty *et al*., 2014, 2015) to sample almost uniformly from ℒ. We call the resulting method <u>T</u>ransm<u>i</u>ssion <u>T</u>ree <u>U</u>niform <u>S</u>ampler (TiTUS).

### 5.3 Consensus Problem

This section introduces a polynomial time algorithm to solve the SCTT problem. The algorithm and the proof for correctness follow the work of (Govek *et al*., 2018). Let 𝒮 = {*S*_1_, …, *S*_*k*_} be a set of *k* transmission trees with edge weights {*w*_1_, …, *w*_*k*_}. Our goal is to find a consensus tree *R* that minimizes *d*(𝒮, *R*) where *d*(·, ·) is the *weighted parent-child distance*. For any given tree *S*_*i*_, we define the function *q*_*i*_ : Σ × Σ → ℕ where

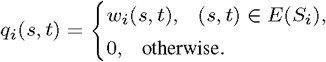

Observe that the parent-child distance between two transmission trees *S*_*i*_ and *S*_*j*_ can be re-written as

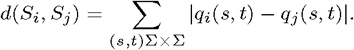

To get the optimal weights for the consensus tree, for any pair of hosts (*s, t*) ∈ Σ × Σ, we define

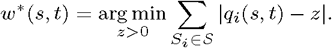

Clearly, *w*\*(*s, t*) for every pair of hosts (*s, t*) is given by max{med, 1} where med is the median of the set {*q*_1_(*s, t*), …, *q*_*k*_(*s, t*)}. Thus, we have the following observation.

**Observation 1.** Given a set 𝒮 = {*S*_1_, …, *S*_*k*_} of *k* transmission trees with edge weights *w*_1_, …, *w*_*k*_, optimal consensus trees *R* that include the edge (*s, t*) must assign this edge weight *w*\*(*s, t*).

We define the *weighted parent-child graph P* as a complete graph with nodes given by the set Σ and a weight function

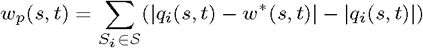

Observe that the weights of the edges of *P* can be negative.

#### Theorem 3.

Given a set 𝒮 = {*S*, …, *S*} of *k* transmission trees with edge weights *w*_1_, …, *w*_*k*_, a minimum weight spanning arborescence of the corresponding weighted parent-child graph *P* defines a tree *R* that is a solution to the SCTT problem with the distance measure used is weighted parent-child distance.

Proof. Provided in Appendix A.4. □

Although edge weights *w*\* of *P* can be negative, the requirement of *R* to be a spanning arborescence of *G* means that we can solve this problem in polynomial time with standard minimum weight spanning arborescence algorithms.

## 6 Results

This section presents the results obtained by applying TiTUS to simulated as well as a real dataset.

### 6.1 Simulations

We employ a two-stage approach to simulate an outbreak, generalizing Didelot *et al*. (2014)’s simulation framework that uses a strong transmission bottleneck to support a weak transmission bottleneck. First, we simulate the transmission process between the *m* hosts using the SIR epidemic model (Allen, 2008). The epidemiological model takes the transmission bottleneck size *κ* and minimum number *n*_*s*_ of strains/leaves for each host *s* as input. Given this input, the model generates a transmission tree *S* with entry *τ*_*e*_(*s*) and removal times *τ*_*r*_ (*s*) for each host *s* as well as the number of transmissions *w*(*s, t*) = *κ* between each pair (*s, t*) ∈ *E*(*S*) of hosts. Given *S* and *w*, we then simulate the evolution of the pathogens within each infected host using a simple coalescence model with constant population size (Kingman, 1982). This process yields a forest of timed phylogenies for each individual host *s*. We construct a single timed phylogeny of all hosts by stitching together individual timed phylogenies using the transmission tree *S*. For each combination of number *m* ∈ {5, 7, 10} of hosts and bottleneck size *κ* ∈ {1, 2, 3} we generate five instances, amounting to a total of 45 simulated instances. The cases with *κ* = 1 correspond to outbreaks with a strong transmission bottleneck. In order to mimic the uncertainty in epidemiological data seen in practice, we increase the length of the entry and removal time interval [*τ*_*e*_(*s*) − Δ, *τ*_*r*_ (*s*) + Δ] for each host *s*, where Δ equals 10% of the total outbreak duration.

We find that increasing the number of hosts and bottleneck size in the simulations leads to an increase in the number of vertices *n* in the phylogenetic trees (Fig. S12a). This leads to a sharp increase in the number of feasible solutions to the rel-DTI (Fig. 3a). The number of solutions to DTI, on the other hand, stays relatively constant for increasing bottleneck size. As a consequence of this, the sampling efficiency of the naive rejection sampling method, defined by the ratio ℒ*/*|ℒ_REL_|, precipitates with increasing number *m* of hosts and bottleneck size *κ* proving it unsuitable for any real applications.

**Fig. 3:**
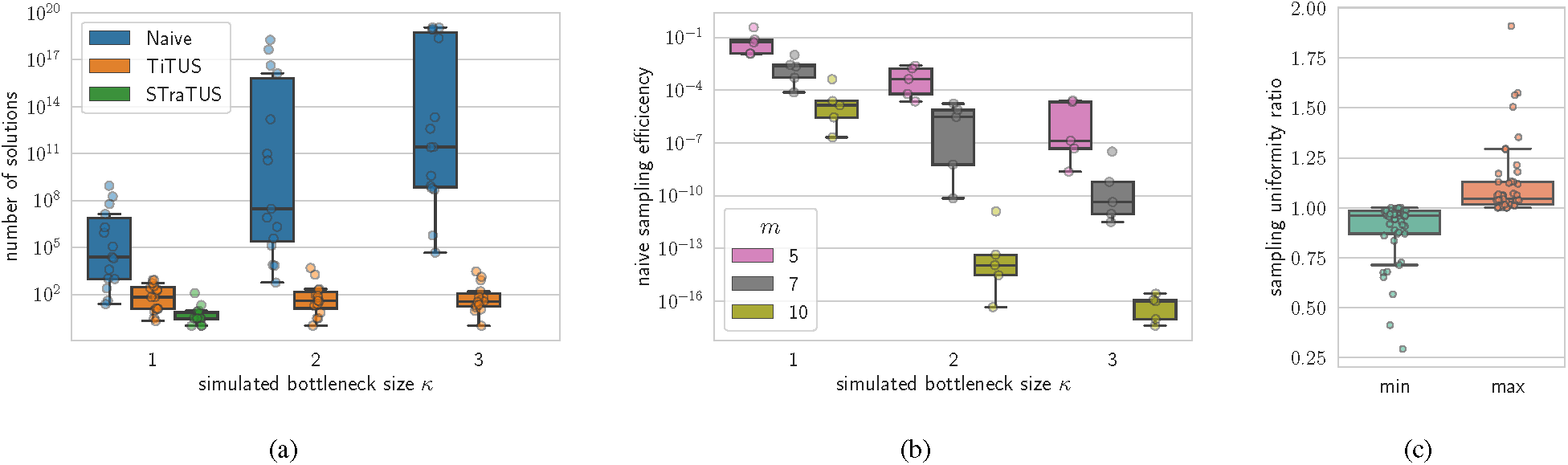
TiTUS accurately samples solutions to the DTI problem. (a) The number of solution to the rel-DTI (|ℒ_REL_|), the DTI (|ℒ|), and the DTI-SB (|ℒ_SB_|) problems computed using the Naive rejection sampling, TiTUS, and STraTUS respectively. The number of solutions to the rel-DTI problem grows rapidly for increasing values of the simulated bottleneck size *κ*, while STraTUS fails to provide any solution when *κ* is greater than 1. (b) The sampling efficiency, defined as the ratio |ℒ| and |ℒ_REL_| for increasing values of simulated number of hosts *m* and bottleneck size *κ*. (c) The ratio between the minimum and maximum observed sampling frequency using TiTUS with the true uniform sampling frequency.

For cases with simulated bottleneck size *κ* > 1, STraTUS fails to provide any solutions (Fig. 3a). This shows that when multi-strain infections occur, transmission history inference with a strong bottleneck assumption will fail to provide the true transmission tree topology. Finally, we assess the sampling accuracy of TiTUS by comparing the sampling frequency with 1*/*|ℒ| where |ℒ| is computed with sharpSAT (Thurley, 2006). For each unique solution that is sampled, the expected sampling frequency 1*/*|ℒ| is the same. Fig. 3c shows that the ratio between both the minimum and maximum values of the observed sampling frequencies with their expected values is close to 1.

In summary, our simulations show that methods that assume a strong transmission bottleneck cannot be applied to outbreaks with a weak bottleneck. Moreover, the exponentially increasing gap between the size of the solution space of rel-DTI compared to DTI renders the rejection-based sampling impractical. In contrast, TiTUS almost uniformly samples from the complex solution space of DTI.

#### 6.1.1 Criteria to Prioritize Candidate Transmission Trees

We propose several criteria for ranking the vertex labelings for a given timed phylogeny uniformly sampled by TiTUS. The first criterion is the *number of transmission edges* in the vertex labeling. Based on the parsimony principle, which has been used in previous works for both phylogeny inference (Sankoff, 1975) as well as transmission tree inference (Wymant *et al*., 2017; Snitkin *et al*., 2012; Sashittal and El-Kebir, 2019), we expect vertex labelings that have few transmission edges to be closer to the ground truth. The second criterion is the *number of unsampled lineages*, which is the number of transmission edges (*u, v*) for which there does not exist a descendant leaf *v*′ (*i*.*e. v* ⪯_*T*_ *v*′) labeled by *ℓ*(*v*). Unsampled lineages are a consequence of multi-strain infections and we expect to see fewer unsampled lineages when the within-host diversity of the infected hosts is adequately sampled. Fig. 5 illustrates this concept.

To assess these criteria, we compare the sampled transmission trees with the ground truth by computing the *infection recall*, defined as the fraction of transmission events between pairs of hosts that are correctly inferred. Fig. 4a shows the value of the *infection recall* for candidate solutions in different percentiles based on the number of transmission edges. Clearly, as we look at solutions with larger transmission numbers, the infection recalls decreases. Fig. 4b show a similar negative correlation between the infection recall and the number of unsampled lineages. We use both the transmission number and the number of unsampled lineages to prioritize the uniformly sampled candidate solutions. Specifically, for any given percentile threshold *α* we include all the vertex labelings whose percentile is at most *α* for both the transmission number and the number of unsampled lineages. (Thus, setting *α* = 1 will include all sampled vertex labelings.) The selected vertex labelings are then used to compute the consensus transmissions tree. Fig. 4c shows the infection recall of the consensus transmission trees for increasing value of the percentile threshold *α*. We see that a value of *α* that is either too small or too large results in a decrease in the *infection recall*. Based on the simulated data, we see that *α*\* = 0.01 yields accurate consensus transmission tree solutions. Hence, the two criteria enable accurate prioritization of sampled vertex labelings.

**Fig. 4:**
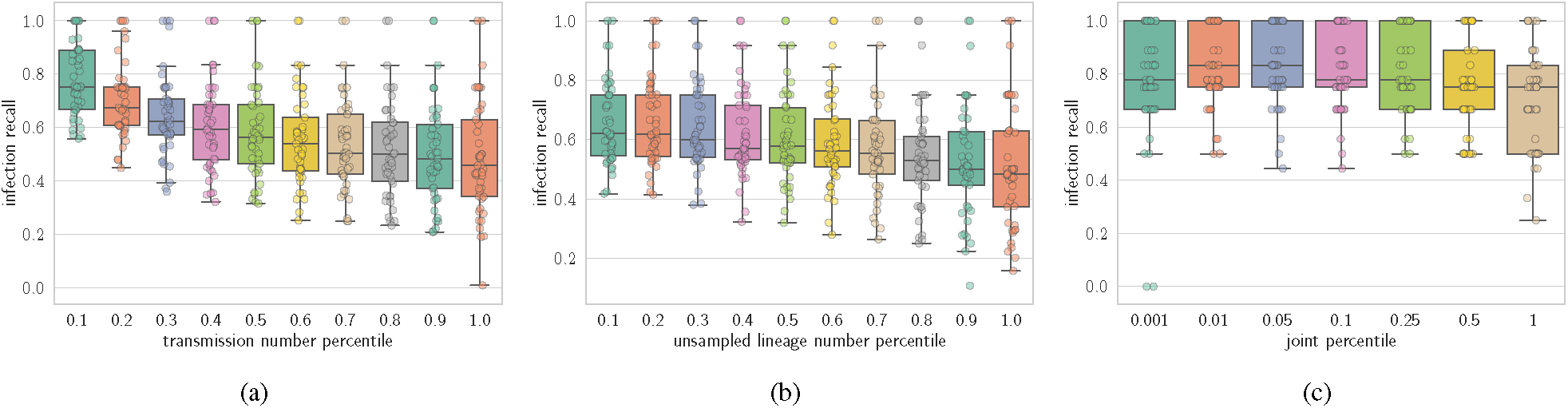
The transmission number and number of unsampled lineages of the solutions to the DTI problem are negatively correlated to the infection recall. (a) The infection recall for the uniformly sampled solution within different percentile based on the transmission number. (b) The infection recall for the uniformly sampled solution within different percentile based on the number of unsampled lineages. (c) The infection recall of the consensus transmission trees within different percentiles of both the transmission number and the number of unsampled lineages simultaneously.

**Fig. 5:**
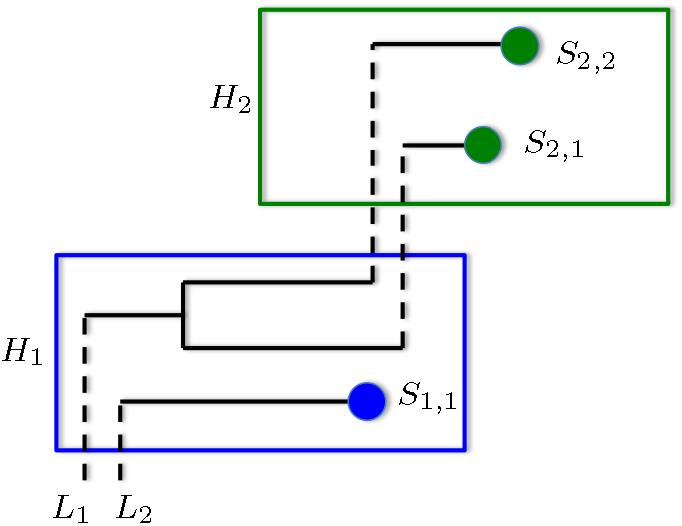
Schematic representation of unsampled lineages in outbreaks. Different hosts *H*_1_ and *H*_2_ are represented by rectangular boxes and the samples taken from the hosts are indicated by blue or green circles inside the boxes respectively. Black lines represent the evolution of pathogen lineages. Solid lines correspond to within-host evolution of the pathogen whereas dashed lines represent the transmission of strains during infection. Two lineages *L*_1_ and *L*_2_ entering host *H*_1_ are shown. Lineage *L*_1_ is an *unsampled lineage* because even though two strains of *L*_1_ are transmitted to host *H*_2_, none of the samples of *H*_1_ belong to the lineage *L*_1_.

### 6.2 HIV Outbreak with a Known Transmission Chain

We apply our method TiTUS to infer the transmission history of an HIV-1 outbreak involving 11 patients with a known transmission chain (Vrancken *et al*., 2014; Lemey *et al*., 2005). The data consists of 212 samples collected over the span of 18 years from the 11 patients. The direction of transmissions and a relatively narrow time interval for each transmission event were inferred from epidemiological information obtained by patient interviews, clinical data and treatment histories of the patients.

The DTI problem for this HIV dataset is set up as follows. For the timed phylogeny, we use the Maximum Clade Credibility (MCC) tree obtained from the partially sequenced *env* regions presented by Vrancken *et al*. (2014) in their publication. Table 1 in Appendix A.6 shows the sampling times and transmission windows provided in the epidemiological data for each of the hosts. The transmission window of a host is the time interval inside of which the host is expected to have been infected. Transmission windows for host A and host D are incongruent with the given timed phylogeny. By this we mean there is no vertex labeling on the given MCC phylogeny that allows for the known transmissions to host A and host D. We exclude these time windows, while the transmission windows for the remaining hosts are used to constraint the possible vertex labelings of the MCC tree. We restrict the infection for each host to take place in within the transmission window provided in the epidemiological data. Appendix A.6 shows the details of the implementation of this constraint in the SAT formulation. Note that while using the time window constraints, we only restrict the time of infection and do not utilize information about the known infectors for each infected host. Finally, for each host the entry time is taken as the beginning of its time window of transmission and the removal time is the latest date of sampling (Table 1). We find that STraTUS fails to provide a solution on this dataset. Indeed, a weak transmission bottleneck needs to be considered in order to infer the transmission history.

**Table 1.** This table shows the epidemiological information provided in the HIV dataset (Vrancken et al., 2014). The transmission window of a host is the expected time-interval during which the host was infected.

| host | transmission window | known infector | latest sample time | entry time | removal time |
| --- | --- | --- | --- | --- | --- |
| A | ? - 14/05/90 | B | 7/11/05 | $\tau(r(T))$ | 7/11/05 |
| F | 01/02/95 - 02/08/95 | A | 19/09/05 | 01/02/95 | 19/09/05 |
| G | 16/01/02 - 16/04/02 | F | 16/04/02 | 16/01/02 | 16/04/02 |
| H | 29/06/95 - 24/07/95 | B | 25/05/98 | 29/06/95 | 25/05/98 |
| I | 01/02/93 - 28/04/93 | B | 06/10/99 | 01/02/93 | 06/10/99 |
| C | 23/09/93 - 10/01/94 | B | 15/12/03 | 23/09/93 | 15/12/03 |
| D | 16/03/95 - 01/07/95 | C | 24/03/03 | 16/03/95 | 24/03/03 |
| L | 23/09/93 - 12/03/06 | C | 24/03/06 | 23/09/93 | 24/03/06 |
| E | 15/06/00 - 01/02/01 | C | 22/02/06 | 15/06/00 | 22/02/06 |
| K | 01/06/04 - 15/09/04 | E | 30/09/04 | 01/06/04 | 30/09/04 |

For this DTI instance, using sharpSAT (Thurley, 2006) we find that there are exactly 30,901,500 feasible vertex labelings. We generate 100,000 samples from this solution space and compute the *infection recall* when compared to the known transmission chain. Fig. 6 shows the values the *infection recall* for solutions with different number of transmission edges and number of unsampled lineages. The infection recall is close to 1 for the solutions that have no unsampled lineages. The number of transmission edges also has a negative, albeit weaker correlation with the infection recall.

**Fig. 6:**
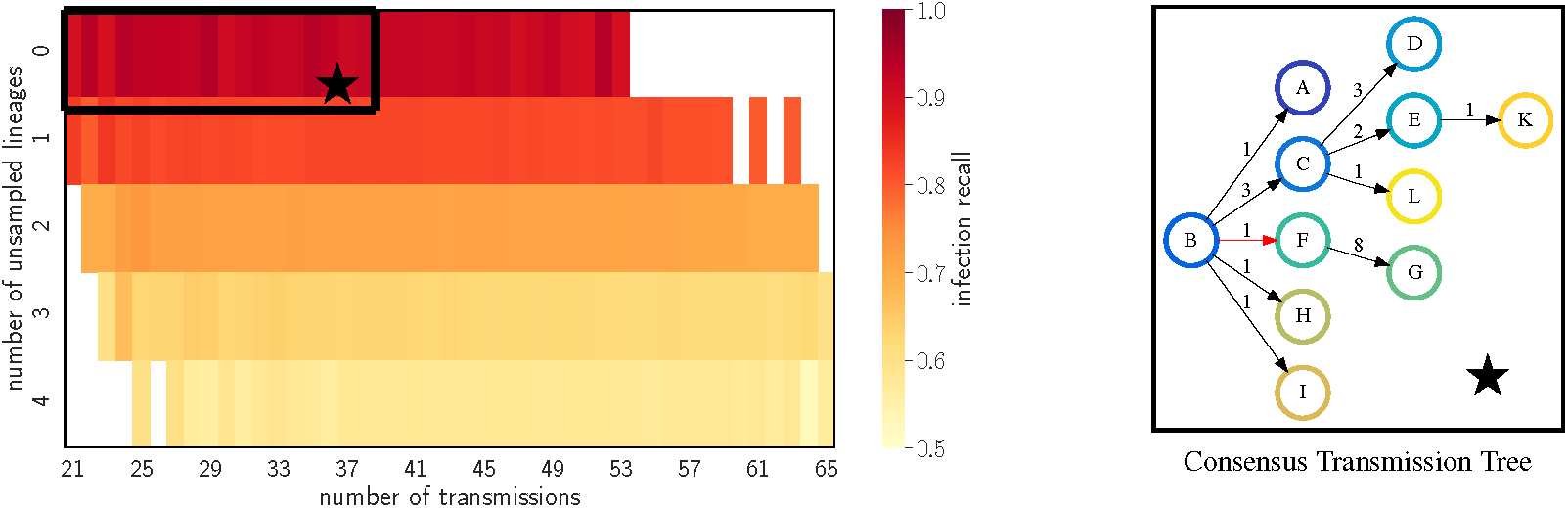
Consensus transmission tree computed for the solutions selected using the proposed criteria infers almost the entire transmission chain for the HIV outbreak. The figure on the left shows the infection recall of the solutions with different transmission numbers and number of unsampled lineages, uniformly sampled using TiTUS. The black box encompasses the solutions selected for the percentile threshold of *α* = 0.01. The figure on the right shows the consensus transmission tree for the selected solutions. Each edge is labeled by the number of strains transmitted from the donor to the recipient host. The incorrectly inferred transmission B→F is highlighted in red.

For any given percentile threshold *α* we include all vertex labelings whose percentile is at most *α* for both the transmission number and the number of unsampled lineages. Based on the simulations, we focus on percentile threshold *α*\* = 0.01. For this threshold value, Fig. 6 shows the consensus transmission tree inferred by TiTUS. The infection recall for this tree is 0.9, *i*.*e*. we correctly infer 9*/*10 transmission from the known transmission chain. We incorrectly infer the transmission B→F while the known transmission to F based on epidemiological data is A→F. Fig. S14 shows similar behavior of the infection recall as a function of *α* as observed in our simulations. Moreover, this figure shows that our method is robust around *α*\* = 0.01.

## 7 Discussion

In this paper, we formulated the Direct Transmission Inference (DTI) problem of inferring transmission trees for a given timed phylogeny and epidemiological data while supporting a weak transmission bottleneck. Weak transmission bottlenecks are common in the spread of diseases due to pathogens with large inoculum sizes, high mutation rates, long incubation times and chronic infections (Leonard *et al*., 2017). Previous studies of counting and sampling transmission trees for a given timed phylogeny assume a strong transmission bottleneck (Kenah *et al*., 2016; Hall and Colijn, 2019), and are not applicable to outbreaks of pathogens with a weak transmission bottleneck, often failing to return any solution.

We proved that the decision version of the DTI problem is NP-complete and the counting version #DTI is #P-complete. Leveraging recent advances made in approximate counting and sampling of solutions to SATISFIABILITY (Chakraborty *et al*., 2014, 2013, 2015; Soos *et al*., 2009), TiTUS, which uses a SATISFIABILITY oracle to almost uniformly sample from the solution space of DTI. In most cases, uniformly sampled candidate solutions from the transmission tree space will deviate considerably from the ground truth. To address this issue, we proposed two criteria that can be used to prioritize the uniformly sampled transmission trees. We demonstrated the performance and robustness of our selection criteria on both simulated data and a real dataset of an HIV outbreak (Vrancken *et al*., 2014).

Further, we also considered the problem of summarizing a given set of candidate transmission tree solutions of a disease outbreak. We defined a new distance metric *weighted parent-child distance* (WPCD) on the space of transmission multi-trees that capture the transmission of multiple strains between hosts during an outbreak. This distance is an extension of the parent-child distance which is used in previous works to summarize cancer phylogenies (Govek *et al*., 2018; Aguse *et al*., 2019). We presented a polynomial time algorithm for finding the consensus transmission tree with minimum total WPCD from the candidate solutions. The performance of the consensus transmission tree of recalling the transmissions that occurred during the outbreak is demonstrated both on simulated and real datasets.

There are several avenues for future research. First, the decision version of the DTI problem can be used to prioritize a posterior distribution of phylogenies, by checking if each phylogeny admits a vertex labeling that induces a transmission tree that is compatible with the given epidemiological data. A similar approach is employed by Sledzieski *et al*. (2019) where they prioritize statistically likely timed phylogenies that admit vertex labelings with fewer transmission edges. By including biological relevant constraints such as a contact map and direct transmission constraints, we expect to obtain high-fidelity phylogenetic and transmission history reconstructions. Second, one limitation of the proposed method is that it assumes that all the infected hosts in the outbreak are sampled. This assumption is only applicable for small outbreaks in regions with perfect surveillance and reporting system in place. An extension of this method to include unsampled hosts would be a useful. Third, akin to (Jombart *et al*., 2017), we plan to extend the SCTT to simultaneously cluster the set 𝒮 of transmission trees and infer a representative consensus transmission tree for each cluster. Finally, we plan to directly include the identified prioritization criteria as constraints in the DTI problem.

---

**Fig. S7:**
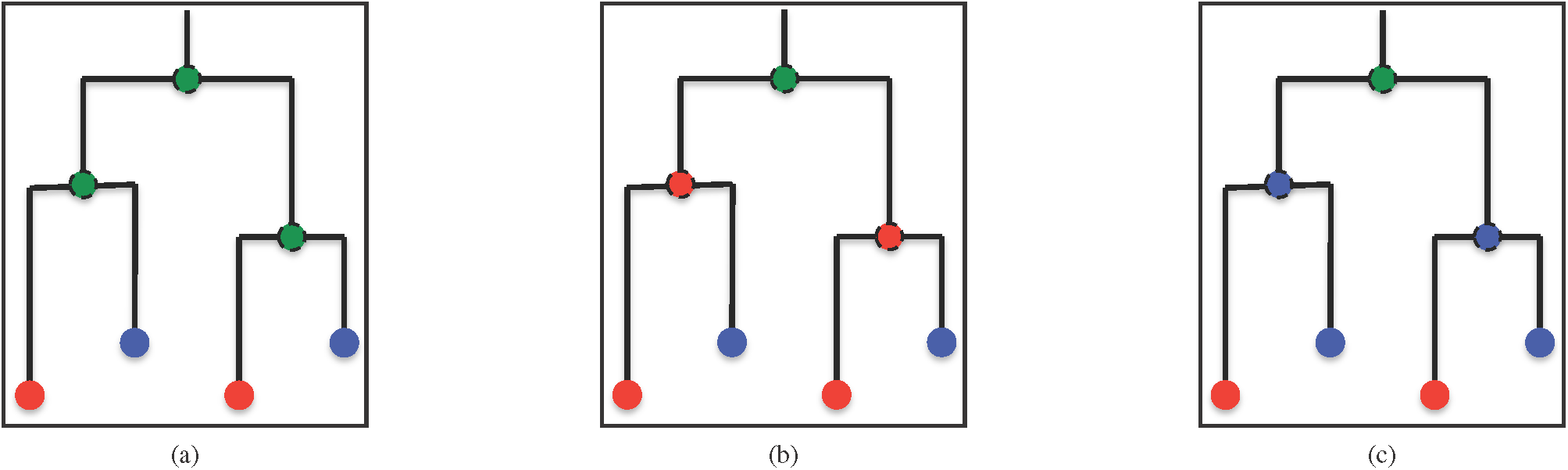
The timed phylogeny shown in Fig. 1a has 3 possible vertex labeling solutions.

## A.1 Background and Theory

In this section we provide the information we could not include in the main text. Fig. S7 shows all the feasible solutions to the representative DTI problem desribed in the Fig. 1.

### A.1.1 Transmission Tree Metric

In this section we show that WPCD is a distance metric. To show that WPCD is a distance metric, for any transmission tree *S*_*i*_, we define the function *q*_*i*_ : Σ × Σ → ℕ as

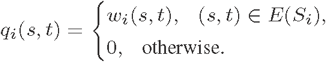

Observe that, by construction, *q*_*i*_ uniquely determines the transmission tree *S*_*i*_ since for any edge (*s, t*) ∈ *E*(*S*_*i*_) we have *w*_*i*_(*s, t*) > 0. Further, the WPCD between any two transmission trees *S*_1_ and *S*_2_ can be alternatively written in terms of *q*_1_ and *q*_2_ as follows,

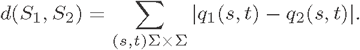

#### Proposition 1.

WPCD is a distance metric on the space of transmission trees 𝒯.

Proof. First, we show that for any two transmission trees *S*_1_ and *S*_2_, *d*(*S*_1_, *S*_2_) = 0 if and only if *S*_1_ = *S*_2_. Clearly when *S*_1_ = *S*_2_, we have *d*(*S*_1_, *S*_2_) = 0. Now, let us consider the case *d*(*S*_1_, *S*_2_) = 0. For any (*s, t*) ∈ Σ × Σ, |*q*_1_(*s, t*) − *q*_2_(*s, t*)| ≥ 0. Therefore, if *d*(*S*_1_, *S*_2_) then for all (*s, t*) ∈ Σ × Σ we have *q*_1_(*s, t*) = *q*_2_(*s, t*) implying that *S*_1_ = *S*_2_.

By definition, WPCD is always nonnegative and symmetric.

We only need to show the triangle inequality, *i*.*e*. given trees *S*_1_, *S*_2_ and *S*_3_, we must show

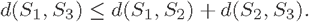

We show this as follows,

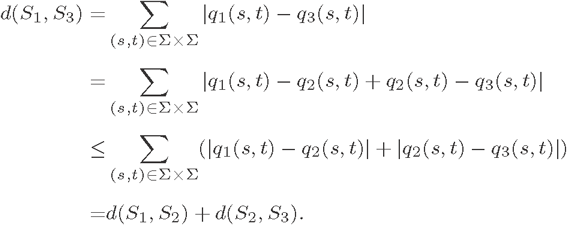

### A.1.2 Sampling Scenarios

The weak transmission bottleneck has some interesting implications for the sampling of the within-host diversity of the infected hosts. Fig. S8 gives an overview, with schematic representations, of 4 different scenarios that can occur for real outbreaks.

**Fig. S8:**
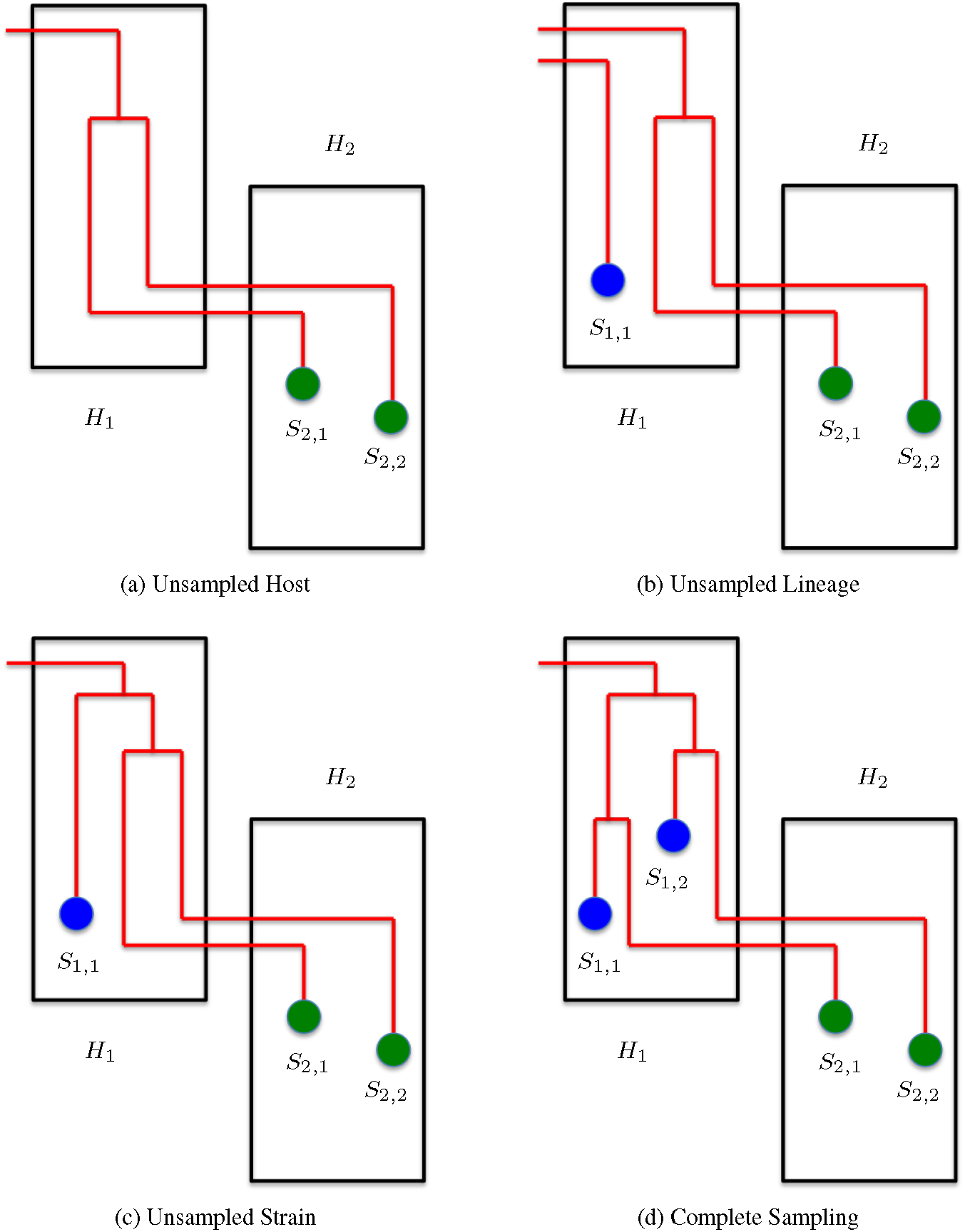
Schematic representation of different sampling scenarios during an outbreak. Different hosts *H*_1_ and *H*_2_ are represented by rectangular boxes and the samples taken from the hosts are indicated by blue or green circles inside the boxes respectively. Red lines represent the evolution of pathogen lineages. Different scenarios described are (a) *Unsampled Host* scenario where host *H*_1_ is not sampled even though it is part of the outbreak and infects *H*_2_ with multiple strains (b) *Unsampled Lineage* where even though host *H*_1_ is sampled with sample *S*_1,1_, the lineage that passes two strains into host *H*_2_ remains unsampled (c) *Unsampled Strain* scenario where the host *H*_1_ is sampled and the right lineage is also sampled however the two strains that are transmitted to host *H*_2_ are not sampled (d) *Complete Sampling* scenario where there is no incomplete lineage sorting (ILS) and all the strains transmitted from *H*_1_ to *H*_2_ are sampled.

## A.2 Complexity

In this section we show the hardness of the decision and the counting versions of the DTI problem using reduction the one-in-three SAT (1-in-3 SAT).

### Problem 4

(1-in-3SAT). Given a Boolean formula 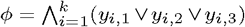 in 3-conjunctive normal form (3-CNF) with *n* variables and *k* clauses, decide whether there exists a truth assignment *θ* : [*n*] → {0, 1} so that each clause has *exactly* one true literal (and thus exactly two false literals).

### A.2.1 Decision Problem

To relate literals to variables, we use the function *ν* : [*k*] × {1, 2, 3} → [*n*] such that *ν*(*i, j*) is the variable corresponding to literal *y*_*i,j*_. We define *σ*(*i, j*) to be 1 if *y*_*i,j*_ is a positive literal (*i*.*e. y*_*i,j*_ = *x*_*ν*(*i,j*)_), otherwise *σ*(*i, j*) = 0 if *y*_*i,j*_ is a negative literal (*i*.*e. y*_*i,j*_ = ¬*x*_*ν*(*i,j*)_). A truth assignment *θ* satisfies *ϕ* if for each clause *i* ∈ [*k*] there exists a *j* ∈ {1, 2, 3} such that *σ*(*i, j*) = *θ*(*ν*(*i, j*)).

**Fig. S9:**
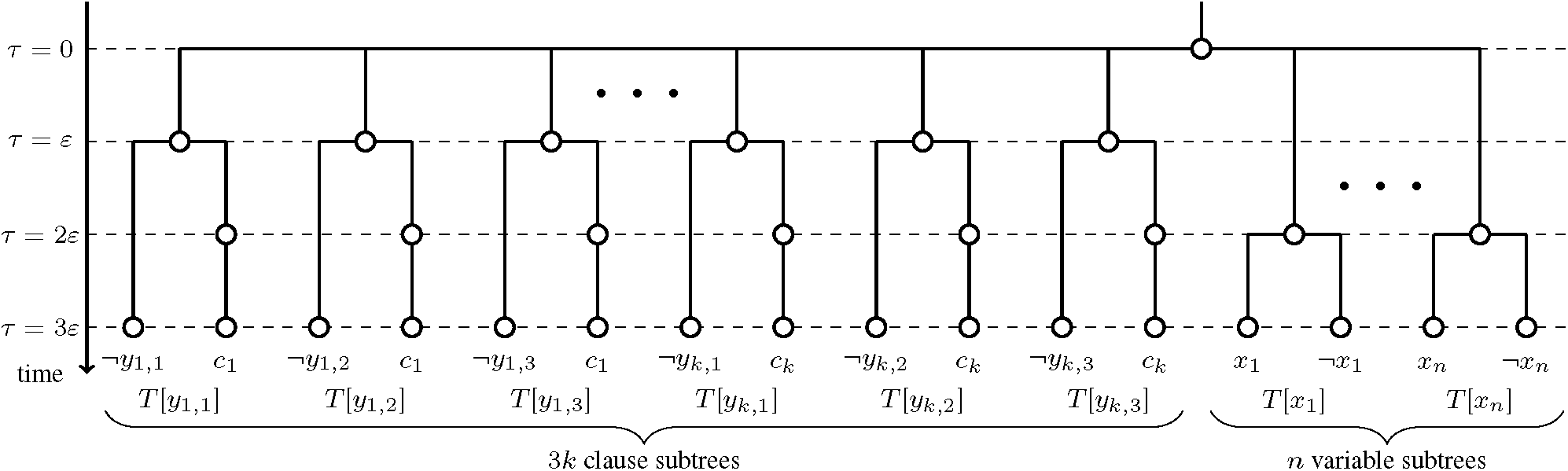
Construction of *T* (*ϕ*) for reduction from 1-in-3SAT to DTI. Let *ϕ* be an 1-in-3SAT formula with *k* clauses and *n* variables. *T* (*ϕ*) is built with a root node *r*(*T* (*ϕ*)) can is connected to 3*k* clause subtrees {*T* [*y*_1,1_], *T* [*y*_1,2_], *T* [*y*_1,3_], …, *T* [*y*_*k*,1_], *T* [*y*_*k*,2_], *T* [*y*_*k*,3_]} and *n* variable subtrees {*T* [*x*_1_], …, *T* [*x*_*n*_]}. We set *τ*_*e*_(⊥) = 0, *τ*_*r*_(⊥) = *ε*, and *τ*_*e*_(*x*_*i*_) = *τ*_*e*_(¬*x*_*i*_) = *ε* and *τ*_*r*_(*x*_*i*_) = *τ*_*r*_(¬*x*_*i*_) = 3*ε* for each variable *i* ∈ [*n*]. For each clause *c*_*i*_, *i* ∈ [*k*] we set *τ*_*e*_(*c*_*i*_) = *τ*_*r*_(*c*_*i*_) = 3*ε*. We prove that there exits a truth assignment so that each clause of *ϕ* has exactly one true literal if and only if there exists a vertex labeling for *T* (*ϕ*) that results in a transmission tree that is a spanning arborescence of the contact map *C*(*ϕ*) (Fig. S10).

**Fig. S10:**
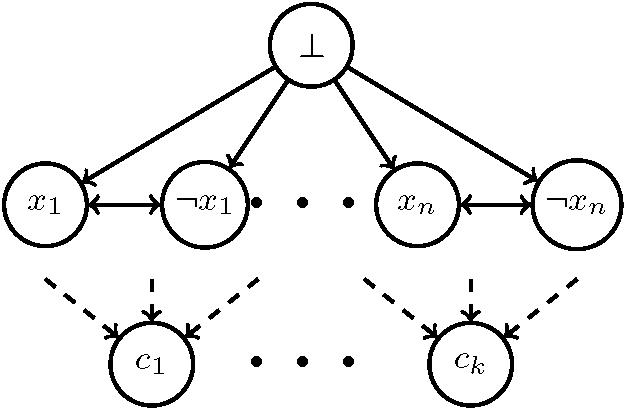
Construction of *C*(*ϕ*) for reduction from 1-in-3SAT to DTI. Let *ϕ* be an 1-in-3SAT formula with *k* clauses and *n* variables. The host set is Σ = {⊥, *x*_1_, …, *x*_*n*_, ¬*x*_1_, …, ¬*x*_*n*_, *c*_1_, …, *c*_*k*_}. We have a directed edge from ⊥ to each of the variables {*x*_1_, …, *x*_*n*_, ¬*x*_1_, …, ¬*x*_*n*_}. Each each *i* ∈ [*n*], variable *x*_*i*_ has an outgoing edge to ¬*x*_*i*_ and similarly variable ¬*x*_*i*_ has an outgoing edge to *x*_*i*_. Finally, each clause *c*_*i*_ has three incoming edges, one from each of the literals that form the clause, i.e. *y*_*i*,1_, *y*_*i*,2_ and *y*_*i*,3_.

Given *ϕ*, we construct a timed phylogeny *T* (*ϕ*) with leaf labeling 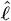, a contact map *C*(*ϕ*) and time-stamps *τ, τ*_*e*_, *τ*_*r*_, as depicted in Fig. S9 and detailed below. We set Σ = {⊥, *x*_1_, …, *x*_*n*_, ¬*x*_1_, …, ¬*x*_*n*_ *c*_1_, …, *c*_*k*_}. Let *ε* > 0 be a small positive constant. As for entry and removal time-stamps, we set *τ*_*e*_(⊥) = 0, *τ*_*r*_(⊥) = *ε*, and *τ*_*e*_(*x*_*i*_) = *τ*_*e*_(¬*x*_*i*_) = *ε* and *τ*_*r*_(*x*_*i*_) = *τ*_*r*_(¬*x*_*i*_) = 3*ε* for each variable *i* ∈ [*n*]. For each clause *c*_*i*_, *i* ∈ [*k*] we set *τ*_*e*_(*c*_*i*_) = *τ*_*r*_(*c*_*i*_) = 3*ε*. Timed phylogeny *T* (*ϕ*) is composed of 3*k* clause gadgets and *n* variable gadgets, each corresponding to a subtree that is directly attached to the root *r*(*T* (*ϕ*)). The root vertex has time-stamp *τ* (*r*(*T* (*ϕ*)) = 0. The leaves of *T* have identical time-stamps 3*ε*. For each variable *i* ∈ [*n*], we have a subtree *T* [*x*_*i*_] whose root has time-stamp *τ* (*r*(*T* [*x*_*i*_])) = 2*ε*. The two children of *r*(*T* [*x*_*i*_]) have identical time-stamps 3*ε*, with one child leading to two leaves labeled by positive literal *x*_*i*_ and the other child leading to two leaves labeled by negative literals ¬*x*_*i*_. Similarly, for each clause *c*_*i*_, *i* ∈ [*k*], we have 3 subtrees *T* [*y*_*i*,1_], *T* [*y*_*i*,2_] and *T* [*y*_*i*,3_]. The root of the subtree *T* [*y*_*i,j*_] has time-stamp *ε* and two children, one of which is the leaf labeled by *x*_*ν*(*i,j*)_ if *y*_*i,j*_ = ¬*x*_*ν*(*i,j*)_ and ¬*x*_*ν*(*i,j*)_ if *y*_*i,j*_ = *x*_*ν*(*i,j*)_. The other child node, denoted as *v*_*i,j*_, has time-stamp *τ* (*v*_*i,j*_) = 2*ε* and has only one child which is a leaf labeled by *c*_*i*_. The contact map *C*(*ϕ*) is constructed as follows. The vertex set for the contact map is given by Σ. We have a directed edge from ⊥ to each of the variables {*x*_1_, …, *x*_*n*_, ¬*x*_1_, …, ¬*x*_*n*_}. For *i* ∈ [*n*], each variable *x*_*i*_ has an outgoing edge to ¬*x*_*i*_ and similarly variable ¬*x*_*i*_ has an outgoing edge to *x*_*i*_. Finally, each clause *c*_*i*_ has three incoming edges, one from each of the literals that form the clause, i.e. *y*_*i*,1_, *y*_*i*,2_ and *y*_*i*,3_. For instance, if *c*_1_ := (*x*_1_ ∨ *x*_2_ ∨ ¬*x*_3_), then we have the directed edges (*x*_1_, *c*_1_), (*x*_2_, *c*_1_) and ¬*x*_3_, *C*_1_. Clearly, *T* (*ϕ*) and *C*(*ϕ*) can be obtained in polynomial time from *ϕ*. An example of this reduction is shown in Fig. S11.

#### Lemma 1.

For any vertex labeling *ℓ* of *T* (*ϕ*), ⊥ is the *root host*.

Proof. Under the direct transmission constraint, *root host* is given by the host that labels the root node of the timed phylogeny. The time stamp of the root node of *T* (*ϕ*) is *τ* (*r*(*T* (*ϕ*))) = 0. The only host that has entry time before *τ*_*e*_ ≤ 0 is ⊥. Therefore, for any vertex labeling we have *ℓ*(*r*(*T* (*ϕ*))) = ⊥, which makes ⊥ the *root host*. □

#### Lemma 2.

For any variable *x*, either {(⊥, *x*), (*x*, ¬*x*)} ⊆ *E*(*S*) or {(⊥, ¬*x*), (¬*x, x*)} ⊆ *E*(*S*).

Proof. For any variable *x*, consider the subtree *T* [*x*]. By construction we have, *τ* (*r*(*T* [*x*])) = 2*ε* and the node only has two children labeled by *x* and ¬*x*. From the contact map we know that the only possible infectors for *x* has ⊥ and ¬*x* and similarly for ¬*x* are ⊥ and *x*. Given that *τ*_*r*_ (⊥) < *τ* (*r*(*T* [*x*])), the only remaining choices for *ℓ*(*r*(*T* [*x*])) are *x* and ¬*x*.

If *ℓ*(*r*(*T* [*x*])) = *x* then we have {(⊥, *x*), (*x*, ¬*x*)} ⊆ *E*(*S*) and if *ℓ*(*r*(*T* [*x*])) = ¬*x* we have {(⊥, ¬*x*), (¬*x, x*)} ⊆ *E*(*S*). □

#### Lemma 3.

For any clause *c*_*i*_ = (*y*_*i*,1_ ∨ *y*_*i*,2_ ∨ *y*_*i*,3_), if (*y*_*i,j*_, *c*_*i*_) ∈ *E*(*S*) then *ℓ*(*r*(*T* [*y*_*i,j*_])) = *y*_*i,j*_ and *ℓ*(*r*(*T* [*y*_*i,j*′_)) = ⊥ for *j*′ ≠ *j*. Proof. Consider the subtree *T* [*y*_*i,j*_]. Let us denote the node that is child of *r*(*T* [*y*_*i,j*_]) and parent of the leaf of *T* [*y*_*i,j*_] labeled with *c*_*i*_ as *v*_*j*_.

**Fig. S11:**
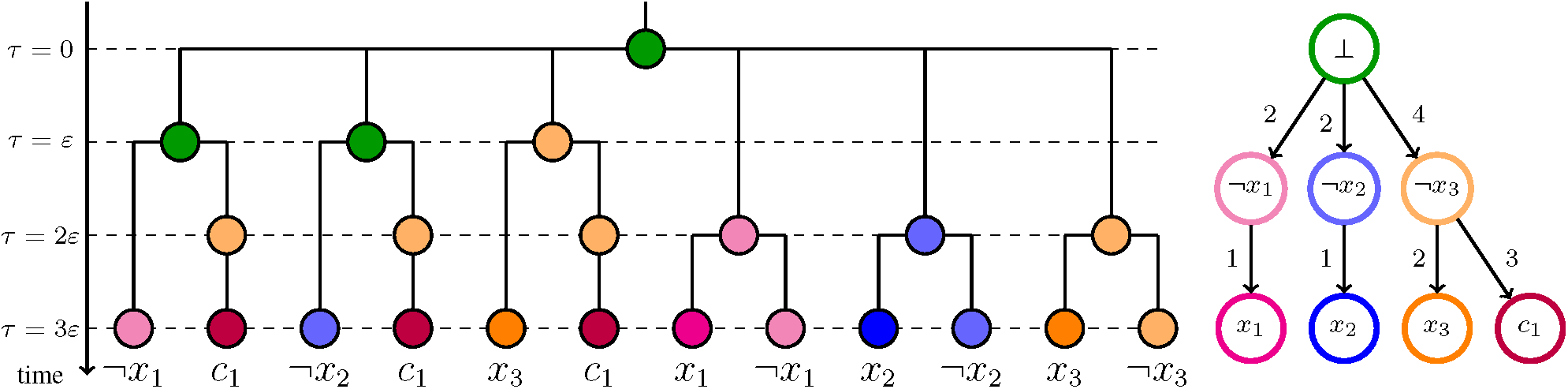
Example of reduction. Consider the 1-in-3SAT Boolean formula *ϕ* = (*x*_1_∨*x*_2_∨¬*x*_3_). *ϕ* is satisfiable with truth assignment *θ*(1) = 0, *θ*(2) = 0 and *θ*(3) = 0. Figures (on the left) shows a vertex labeling *ℓ* corresponding to *θ*. Since the vertex labeling admits a transmission tree (one the right), *ϕ* is Exactly-1 satisfied with truth assignment *θ*.

Since *S* is a spanning arborescence of *C*(*ϕ*) we have either (*y*_*i*,1_, *c*_*i*_), (*y*_*i*,2_, *c*_*i*_) or (*y*_*i*,3_, *c*_*i*_) in *E*(*S*). Without loss of generality, let us assume that (*y*_*i*,1_, *c*_*i*_) ∈ *E*(*S*).

The edges (*v*_1_, *δ*_*T*_ (*v*_1_)), (*v*_2_, *δ*_*T*_ (*v*_2_)) and (*v*_3_, *δ*_*T*_ (*v*_3_)) need to be transmission edges since *τ* (*v*_1_) = *τ* (*v*_2_) = *τ* (*v*_3_) < *τ*_*e*_(*c*_*i*_). Since (*y*_*i*,1_, *c*_*i*_) ∈ *E*(*S*), we require *ℓ*(*v*_1_) = *ℓ*(*v*_2_) = *ℓ*(*v*_3_) = *y*_*i*,1_. Looking at *r*(*T* [*y*_*i*,2_]) and *r*(*T* [*y*_*i*,3_]), since each clause consists of distinct variables, we can only have *ℓ*(*r*(*T* [*y*_*i*,2_])) = *ℓ*(*r*(*T* [*y*_*i*,3_])) = ⊥. Consequently, the transmission edges (*r*(*T* [*y*_*i*,2_]), *v*_2_) and (*r*(*T* [*y*_*i*,3_]), *v*_3_) results in a edge (⊥, *y*_*i*,1_) in *E*(*S*). By Lemma 2, this also means (*y*_*i*,1_, ¬*y*_*i*,1_) ∈ *E*(*S*) and therefore *ℓ*(*r*(*T* [*y*_*i*,1_])) = *y*_*i*,1_. □

#### Lemma 4.

For any literal *y*_*i,j*_ in clause *c*_*i*_, (⊥, *y*_*i,j*_) ∈ *E*(*S*) if and only if (*y*_*i,j*_, *c*_*i*_) ∈ *E*(*S*).

Proof. Consider the subtree *T* [*y*_*i,j*_]. Let us denote the node that is child of *r*(*T* [*y*_*i,j*_]) and parent of the leaf of *T* [*y*_*i,j*_] labeled with *c*_*i*_ as *v*.

(⇒) If (⊥, *y*_*i,j*_) ∈ *E*(*S*), then by Lemma 2 we know that (*y*_*i,j*_, ¬*y*_*i,j*_) ∈ *E*(*S*). Therefore, *ℓ*(*r*(*T* [*y*_*i,j*_])) = *y*_*i,j*_. Given that *ℓ*(*r*(*T* [*y*_*i,j*_])) = *y*_*i,j*_, *ℓ*(*δ*_*T*_ (*v*)) = *c*_*i*_ and *τ* (*v*) = *ε*, the only feasible label for *v* is *y*_*i,j*_. Therefore *ℓ*(*v*) = *y*_*i,j*_ and (*y*_*i,j*_, *c*_*i*_) ∈ *E*(*s*).

(⇐) If (*y*_*i,j*_, *c*_*i*_) ∈ *E*(*S*), then since *τ* (*v*) < *τ*_*e*_(*c*_*i*_), we have *ℓ*(*v*) = *y*_*i,j*_. From Lemma 3 we know that *ℓ*(*r*(*T* [*y*_*i,j*_])) is either ⊥ or *y*_*i,j*_. If *ℓ*(*r*(*T* [*y*_*i,j*_])) = ⊥, then we will have {(⊥, *y*_*i,j*_), (⊥, ¬*y*_*i,j*_)} which is not possible due to Lemma 2. Therefore *ℓ*(*r*(*T* [*y*_*i,j*_])) = *y*_*i,j*_ and consequently (⊥, *y*_*i,j*_) ∈ *E*(*S*). □

#### Proposition 2.

There exists a vertex labeling *ℓ* of *T* (*ϕ*) under the direct transmission constraint such that the corresponding transmission tree *S*(*ℓ*) is a spanning arborescence of *C*(*ϕ*) if and only if *ϕ* is satisfiable with a truth assignment *θ* so that each clause has exactly one true literal.

Proof. (⇒) Let *ℓ* be a vertex labeling of *T* (*ϕ*) under the direct transmission constraint such that the corresponding transmission tree *S* is a spanning arborescence of *C*(*ϕ*). We construct the corresponding truth assignment *θ* for *ϕ* as follows. From Lemma 2 we know that for any variable *x*, either (⊥, *x*) ∈ *E*(*S*) or (⊥, ¬*x*) ∈ *E*(*S*). We set *θ*(*i*) = 1 if (⊥, *x*_*i*_) ∈ *E*(*S*) and *θ*(*i*) = 0 if (⊥, ¬*x*_*i*_) ∈ *E*(*S*). We claim that the this truth assignment satisfies *ϕ* with exactly one literal for each clause.

We need to show that, for any clause *c*_*i*_ = (*y*_*i*,1_ ∨ *y*_*i*,2_ ∨ *y*_*i*,3_), exactly one of (⊥, *y*_*i*,1_), (⊥, *y*_*i*,2_) and (⊥, *y*_*i*,3_) is in *E*(*S*). From Lemma 4 we know that (⊥, *y*_*i,j*_) ∈ *E*(*S*) if and only if (*y*_*i,j*_, *c*_*i*_) ∈ *E*(*S*). Since *S* is a spanning arborescence, exactly one of (*y*(*i*, 1), *c*_*i*_), (*y*_*i*,2_, *c*_*i*_) and (*y*_*i*,3_, *c*_*i*_) is in *E*(*S*). Therefore, exactly one of (⊥, *y*_*i*,1_), (⊥, *y*_*i*,2_) and (⊥, *y*_*i*,3_) is in *E*(*S*) which renders the clause *c*_*i*_ satisfied with exactly one literal.

(⇐) Consider the truth assignment *θ* that satisfies *ϕ* with exactly one literal for each clause in *ϕ*. We build the vertex labeling *ℓ* for *T* (*ϕ*) as follows. From Lemma 1 it is clear that ⊥ is the root host and therefore *r*(*S*) = ⊥. We set *ℓ*(*T* [*x*_*i*_]) = *x*_*i*_ if *θ*(*i*) = 1 and *ℓ*(*T* [*x*_*i*_]) = ¬*x*_*i*_ if *θ*(*i*) = 0. For any clause *c*_*i*_ in *ϕ*, if *y*_*i,j*_ is true we set *ℓ*(*r*(*T* [*y*_*i,j*_])) = *y*_*i,j*_ and if ¬*y*_*i,j*_ is true we set *ℓ*(*r*(*T* [*y*_*i,j*_])) = ⊥. Finally, we set *ℓ*(*v*_*i,j*_) = *y*_*i,j*_ for all *j* ∈ {1, 2, 3}. We need to show that constructed vertex labeling satisfies the direct transmission constraint and that the resulting transmission tree is a spanning arborescence of the contact map *C*(*ϕ*). We do this by first showing that (i) each variable has a unique infector and (ii) all transmission edges between the same pair of hosts have time intervals that overlap.

Consider all the variables that are assigned true by the truth assignment. The infector for all these variables is ⊥ since *ℓ*(*r*(*T* (*ϕ*))) = ⊥ and *ℓ*(*T* [*x*_*i*_]) = *x*_*i*_ if *θ* = 1 and *ℓ*(*r*(*T* [*y*_*i,j*_])) = ⊥ if ¬*y*_*i,j*_ is true. This agrees with *C*(*ϕ*). The time intervals of the outgoing edges from *r*(*T* (*ϕ*)) and *r*(*T* [*y*_*i,j*_]), ∀*i* ∈ [*k*], *j* ∈ {1, 2, 3} contain *τ* = *ε*. Therefore, all possible transmission edges from ⊥ overlap at *τ* = *ε*.

Consider the variables that are assigned false by the truth assignment. From Lemma 2 we know that for any such variable *x*, they are infected by ¬*x*. This agrees with *C*(*ϕ*). Moreover, these variables do not label any of the interval vertices of the tree *T* and all the leaves of *T* are at the same time-stamp *τ* = 3*ε*. Therefore, all possible transmission edges to any such variable *x* overlap at *τ* = 3*ε*.

Finally, consider any clause *c*_*i*_. All the internal vertices *v*_*i,j*_, *j* ∈ {1, 2, 3} are labeled by the same variable *y*_*i,j*_ that renders the clause *c*_*i*_ satisfied. As a result, *y*_*i,j*_ is a unique infector of *c*_*i*_ and (*y*_*i,j*_, *c*) exists in *E*(*C*(*ϕ*)) by construction. Also, time-stamp of all vertices *v*_*i,j*_ are the same *τ* = 2*ε* and therefore, the transmission edges overlap at *τ* = 2*ε*. □

### A.2.2 Counting Problem

This section proves the #P-completeness of the #DTI problem.

#### Proposition 3.

There exists a parsimonious reduction from #1-in-3SAT to #DTI.

Proof. Consider the reduction shown in Section 4. Here we show that this reduction is parsimonious, i.e. it preserves the number of solutions in the solution spaces of the two problems. We show a bijection between the solution space of a 1-in-3SAT and the solution space of the corresponding DTI instance.

Consider the Boolean formula *ϕ*. For a given truth assignment *θ* that satisfies each clause of *ϕ* with exactly one true literal, we construct the vertex labeling of *T* (*ϕ*) as following. We let *ℓ*(*T* [*x*_*i*_]) = *x*_*i*_ if *θ*(*i*) = 1 and *ℓ*(*T* [*x*_*i*_]) = ¬*x*_*i*_ if *θ*(*i*) = 0. We will show that this unique determines the labeling for the rest of the internal vertices of *T* (*ϕ*). Consider the clause *c*_*i*_ and the corresponding subtrees *T* [*y*_*i*,1_], *T* [*y*_*i*,2_] and *T* [*y*_*i*,3_]. Since the truth assignment satisfies each clause with exactly one literal, without loss generality, assume that *y*_*i*,1_ is true. Then using Lemma 4, since (⊥, *y*_*i,j*_) ∈ *E*(*S*), we have (*y*_*i,j*_, *c*_*i*_) ∈ *E*(*S*). For the nodes *v*_*i,j*_ we have *τ* (*v*_*i,j*_) < *τ*_*e*_(*c*_*i*_) and therefore *ℓ*(*v*_*i,j*_) = *y*_*i,j*_, ∀*j* ∈ {1, 2, 3}. Finally, the vertex labels for the roots of the clause subtrees *ℓ*(*r*(*T* [*y*_*i*,1_])) = *ℓ*(*r*(*T* [*y*_*i*,2_])) = *ℓ*(*r*(*T* [*y*_*i*,3_])) = *y*_*i*,1_ due to Lemma 3. Proof of Proposition 2 shows that this vertex labeling is a solution of the DTI problem.

From a given vertex labeling *ℓ*, we construct the truth assignment as follows. We set *θ*(*i*) = 1 if *ℓ*(*r*(*T* [*x*_*i*_])) = *x*_*i*_ and *θ*(*i*) = 0 if *ℓ*(*r*(*T* [*x*_*i*_])) = ¬*x*_*i*_. Proof of Proposition 2 shows that this is a truth assignment that satisfies each clause with exactly one true literal.

The construction of *θ* from *ℓ* and *ℓ* from *θ* are inverses of each other. If we view these constructions as functions then they show a bijection in the solutions spaces of #1-in-3SAT and #DTI. This shows that the number of solutions is preserved. Obviously, the reduction can be performed in polynomial time. Therefore, the reduction is parsimonious. □

## A.3 Naive Rejection Sampling Algorithm

Here we descirbe the naive rejection sampling algorithm introduced in Section 5.2.1. Let *h*[*v, s*] denote the number of vertex labelings *ℓ* ∈ ℒ_REL_ in the subtree *T*_*v*_ of *T* rooted at vertex *v* when *ℓ*(*v*) = *s*. We define *h*[*v, s*] recursively as

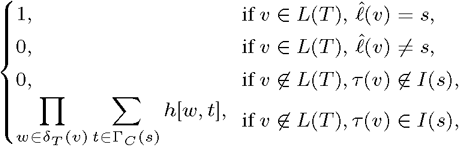

where *I*(*s*) = [*τ*_*e*_(*s*), *τ*_*r*_(*s*)] and G_*C*_ (*s*) = {*s, δ*_*C*_ (*s*)}. Let Σ* = {*s*_1_, …, *s*_*k*_} be the set of possible labels for the root vertex *r*(*T*), *i*.*e*. Σ* = {*s* ∈ Σ | *τ* (*r*(*T*)) ∈ *I*(*s*)}. The number of vertex labelings |ℒ_REL_| is given by Σ_*s*′∈Σ*_ *h*[*r*(*T*), *s*′].

Using the count matrix *h*[*u, s*], we introduce a subroutine that takes a vertex *v* and host *s* as input, and uniformly samples a vertex labeling *ℓ*_*u*_ of subtree *T*_*u*_ rooted at *u* subject to the restriction that *ℓ*_*u*_(*u*) = *s* (Algorithm 3). The fraction *p*_*s*_ of the vertex labelings *ℓ* where *ℓ*(*r*(*T*)) = *s* equals *h*[*r*(*T*), *s*]/Σ_*s*′∈Σ*_ *h*[*r*(*T*), *s*′]. Thus, to sample *all* vertex labelings uniformly at random, we draw a *s* ∈ Σ* according to the categorical probability distribution defined by (*p*_1_, …, *p*_*k*_). Algorithm 4 is then used on *T* with *ℓ*(*r*(*T*)) = *s* to sample minimum transmission host labeling *ℓ* of *T* uniformly at random. This takes *O*(*nm*) time per sample.

For a given phylogeny and vertex labeling (*T, ℓ*), it is possible to find the minimum number of transmission events in polynomial time (Sashittal and El-Kebir, 2019). The *direct transmission constraint* is satisfied by the vertex labeling when the number of transmission events is *m* − 1, where each transmission event corresponds to an edge of the transmission tree. We can therefore draw vertex labelings from ℒ_REL_ and only retain the solutions that belong to ℒ in polynomial time. Since we are uniformly sampling from ℒ_REL_, the retained solutions will also be uniformly sampled from ℒ. For the counting problem we estimate the number of vertex labelings in ℒ by the success rate of the sampling algorithm. Say after *K* draws of samples from ℒ_REL_, we retain *K*′ vertex labelings that belongs to ℒ. In that case the estimate of the size of ℒ, denote by ⟨|ℒ|⟩, is given by

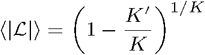

From the law of large numbers, as *K* → ∞ we have ⟨|ℒ|⟩ → |ℒ|. We now present the algorithms for naive rejection based sampling.

### Algorithm 1 EnumRelDTI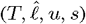

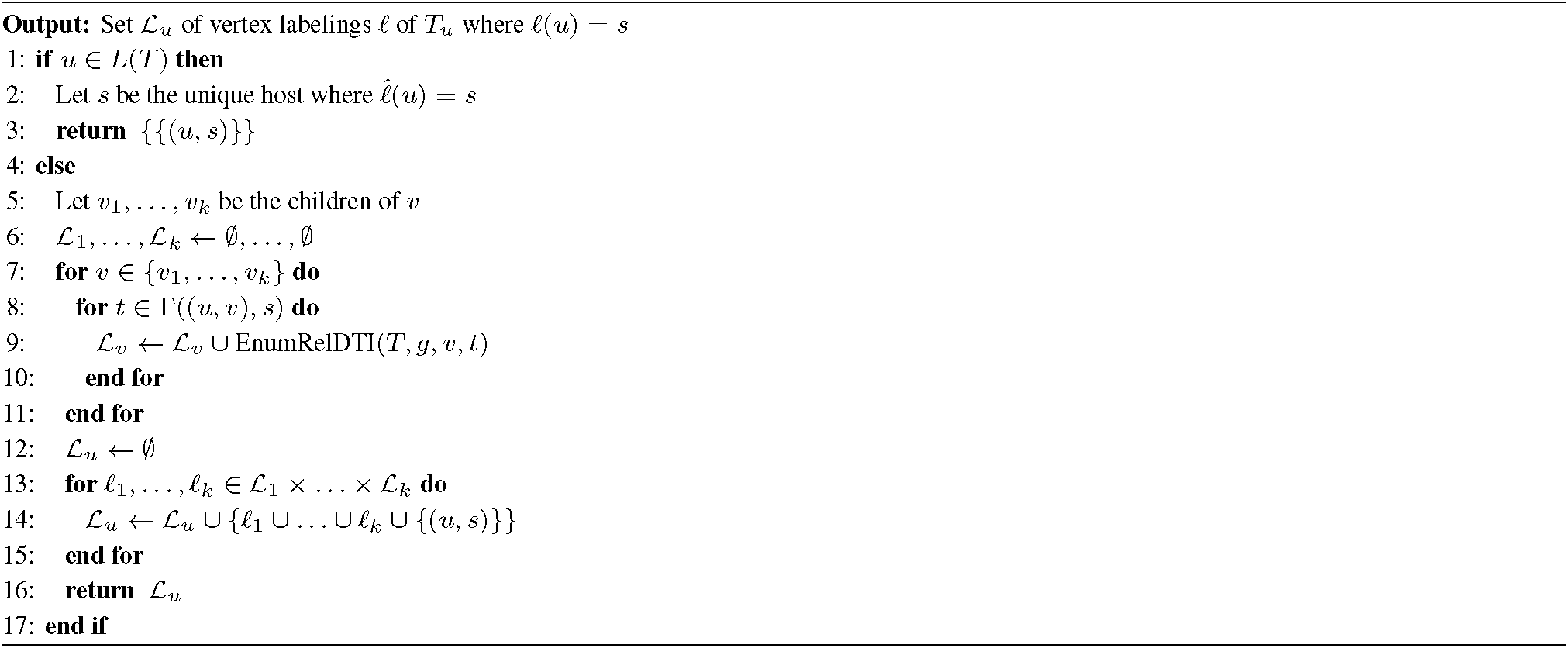

### Algorithm 2 EnumRelDTI(*T, g*)

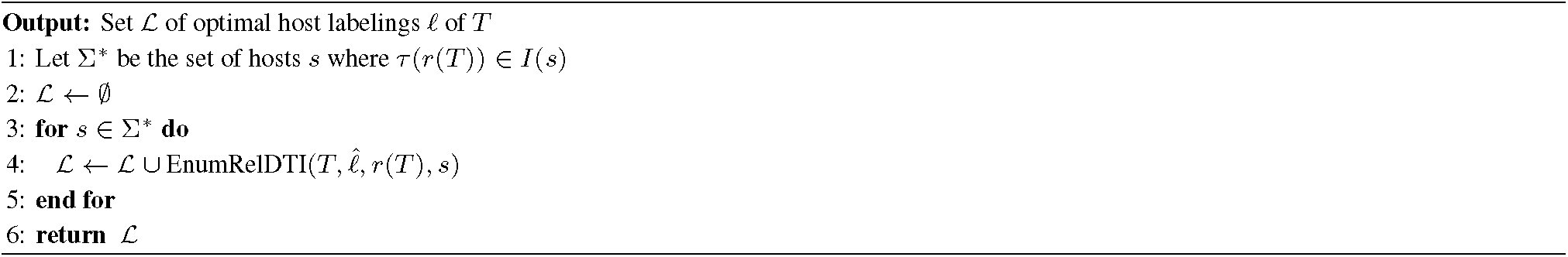

### Algorithm 3 SampleRelDTI(*T, h, u, s*)

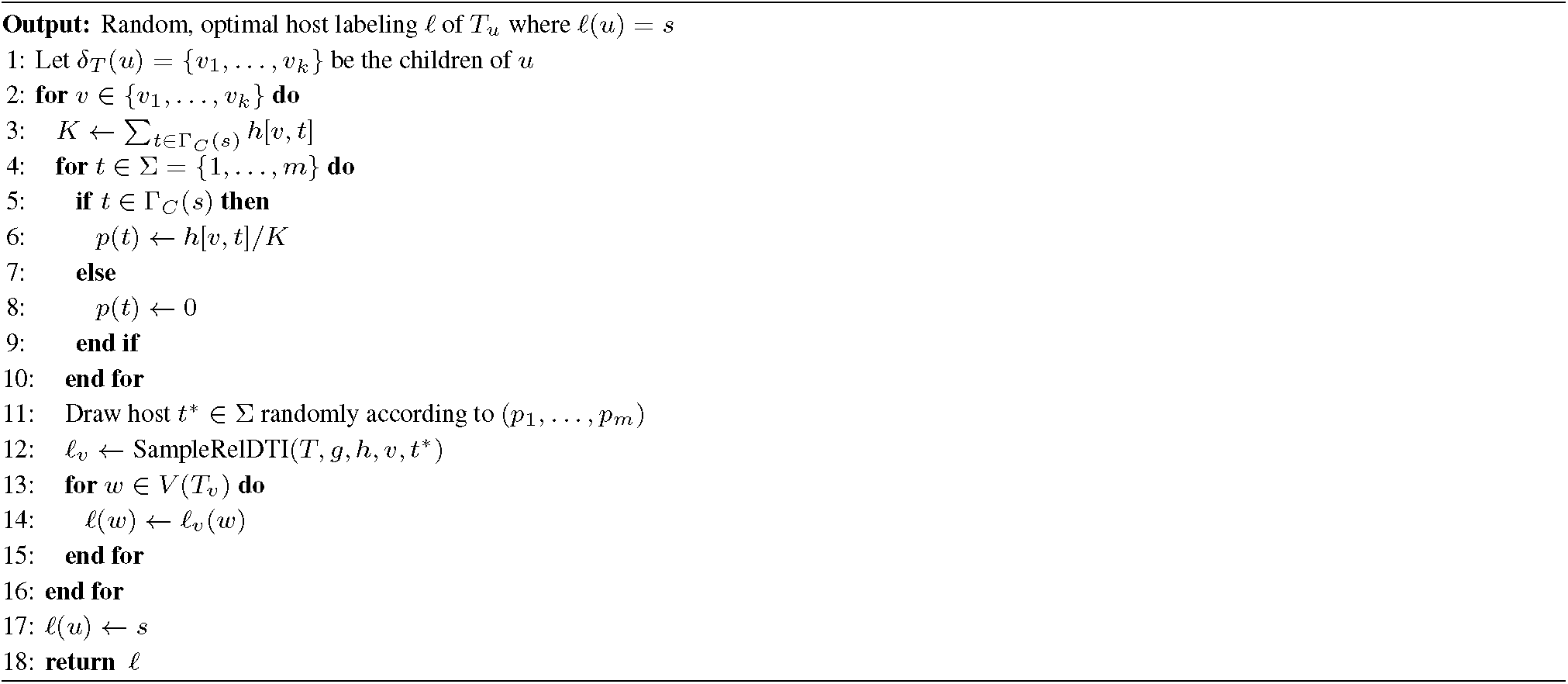

### Algorithm 4 SampleRelDTI(*T, h*)

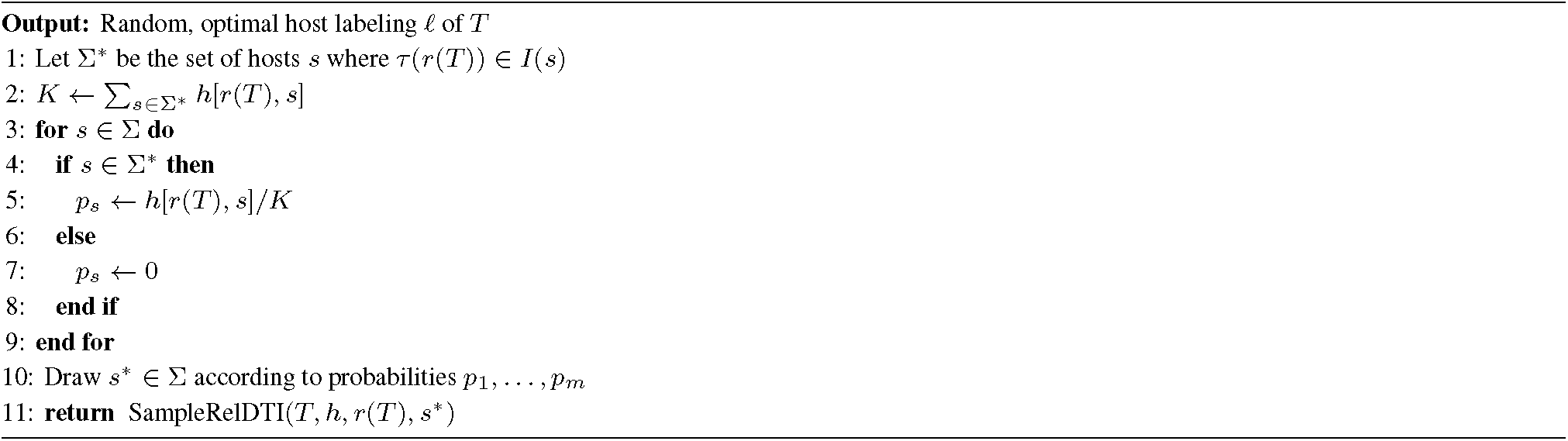

## A.4 Consensus Transmission Tree Algorithm Proof

### Theorem 4.

Given a set 𝒮 = {*S*_1_, …, *S*_*k*_} of *k* transmission trees with edge weights 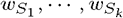, the minimum weight spanning arborescence of the corresponding weighted parent-child graph *P* defines a tree *R* that is a solution to the SCTT problem with the distance measure used is weighted parent-child distance.

Proof. Consider the *weighted parent-child graph P* for the set of transmission trees 𝒮. Since *P* is a complete graph, the optimal consensus tree *R* is necessarily a spanning arborescence of *P*. The weights of the edges in *R* are given by *w*\* due to Proposition 1. The total WPCD of *R* from the set of transmission trees 𝒮 is given by 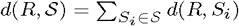 where

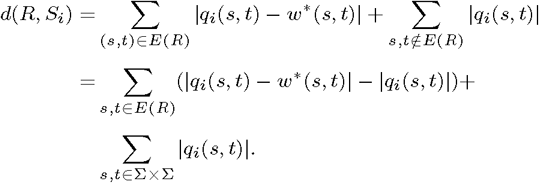

Consequently,

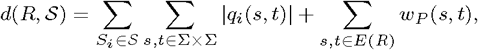

where the first term is a constant with respect to *R* and minimizing the second term is equivalent to finding the minimum weight spanning arborescence of *P*. □

## A.5 Additional Simulation Results

**Fig. S12:**
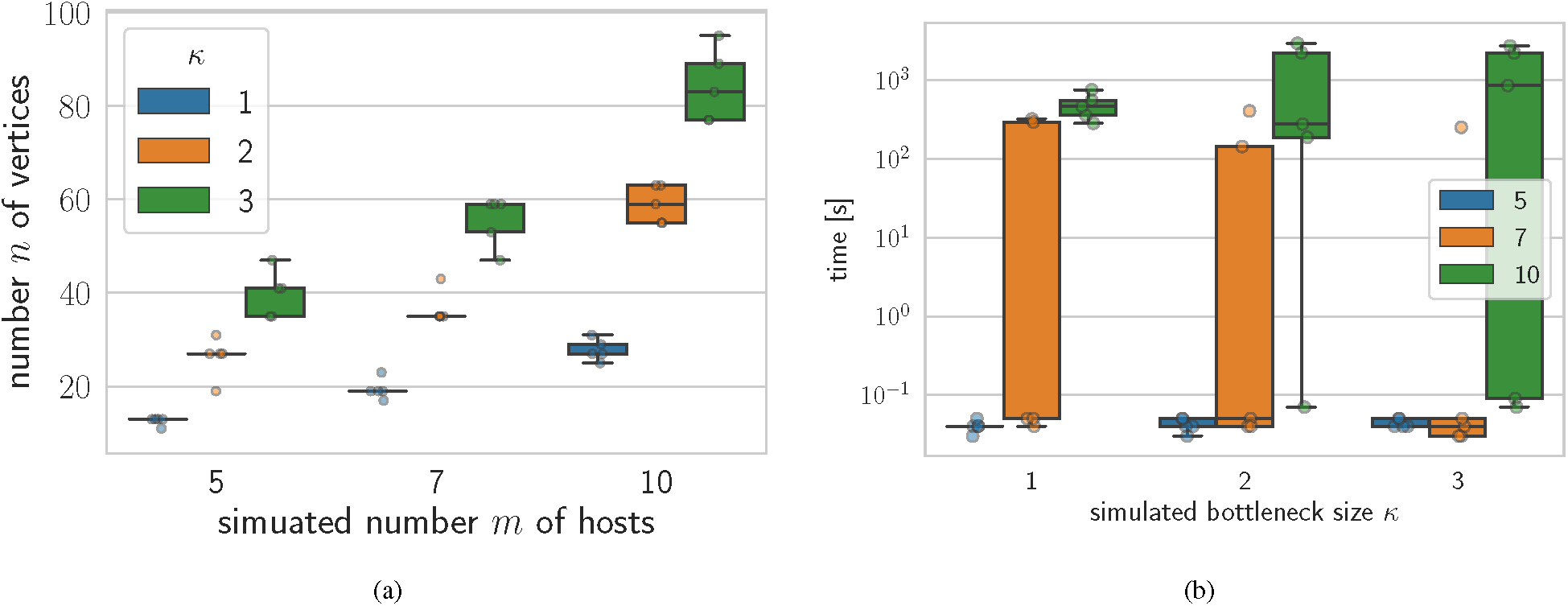
(a) The number of vertices *n* in the timed phylogeny *T* for increasing number *m* of simulated hosts and bottleneck size *κ*. (b) Time taken to generate 100,000 uniformly sampled solutions to the DTI problem using TiTUS for increasing values of simulated bottleneck size *κ*.

## A.6 Additional HIV Data Analysis and Implementation Details

**Fig. S13:**
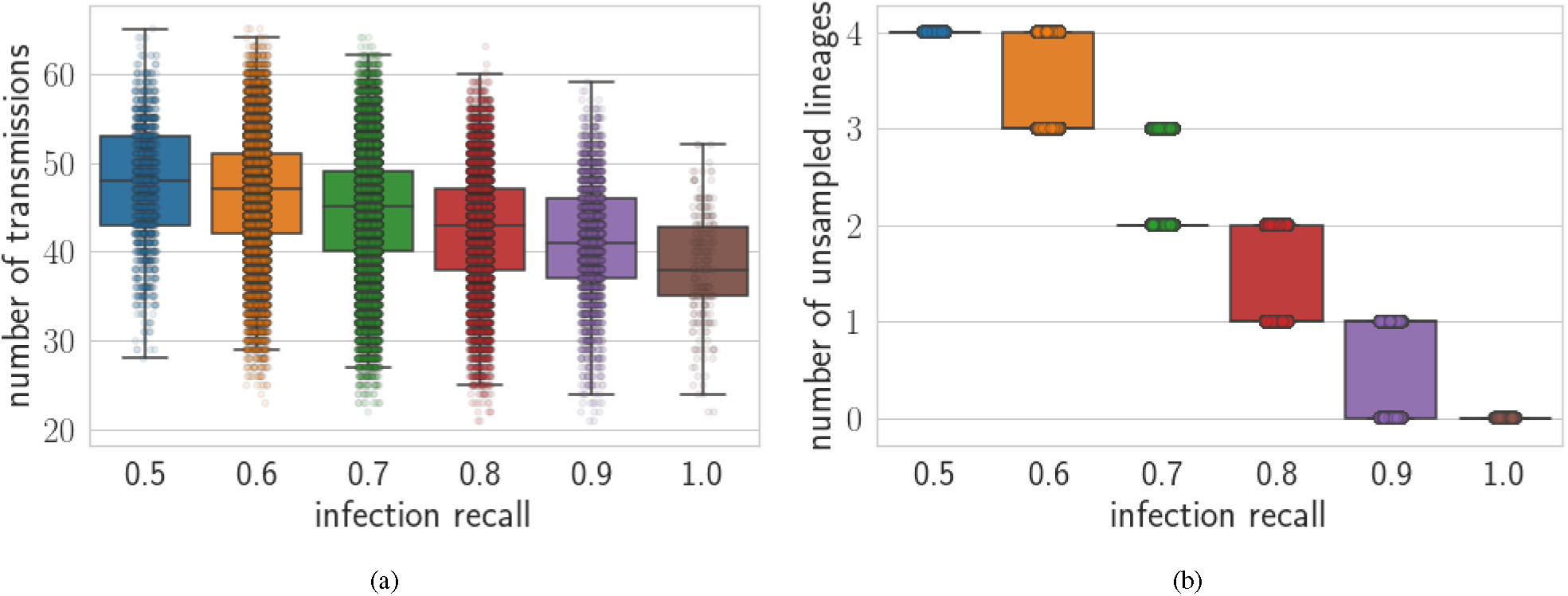
(a) Transmission number and (b) number of unsampled lineages of all the solutions generated using TiTUS on the HIV dataset vs different infection recall values.

**Fig. S14:**
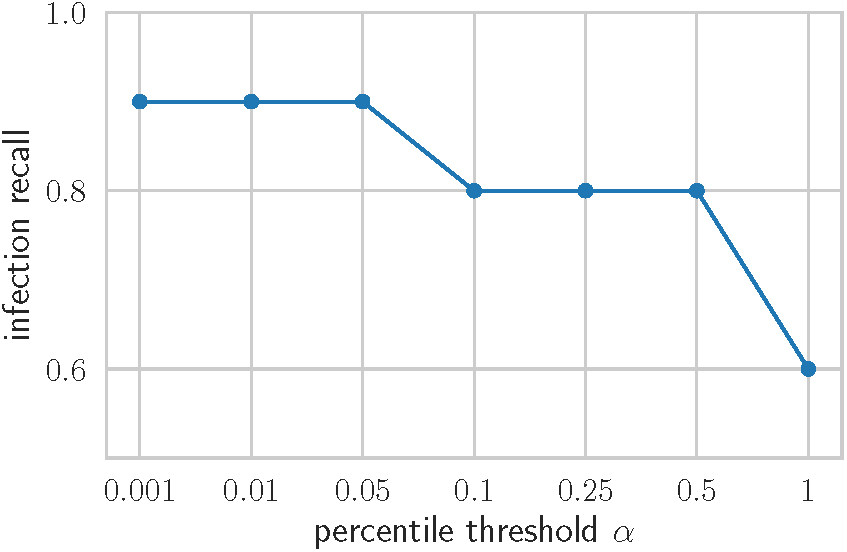
The infection recall of the consensus transmission tree for solutions sampled using TiTUS on the HIV dataset for increasing values of the percentile threshold *α*.

